# Microbial interkingdom interactions in roots promote Arabidopsis survival

**DOI:** 10.1101/354167

**Authors:** Paloma Durán, Thorsten Thiergart, Ruben Garrido-Oter, Matthew Agler, Eric Kemen, Paul Schulze-Lefert, Stéphane Hacquard

**Affiliations:** Max Planck Institute for Plant Breeding Research, 50829 Cologne, Germany.; Cluster of Excellence on Plant Sciences (CEPLAS), Max Planck Institute for Plant Breeding Research, 50829 Cologne, Germany.

**Author notes:** Institute of Microbiology, Friedrich Schiller University, 07743 Jena, Germany (Matthew Agler). Institute of Plant Biochemistry, ZMBP, University of Tübingen, 72076 Tübingen, Germany. Co-first author (Paloma Durán, Thorsten Thiergart). Co-senior author (Paul Schulze-Lefert, Stéphane Hacquard). Correspondence (S.H.), (P.S.-L.).

## Abstract

Roots of healthy plants are inhabited by soil-derived bacteria, fungi, and oomycetes that have evolved independently in distinct kingdoms of life. How these microorganisms interact and to what extent those interactions affect plant health are poorly understood. We examined root-associated microbial communities from three *Arabidopsis thaliana* populations and detected mostly negative correlations between bacteria and filamentous microbial eukaryotes. We established microbial culture collections for reconstitution experiments using germ-free *A. thaliana.* In plants inoculated with mono- or multi-kingdom synthetic microbial consortia, we observed a profound impact of the bacterial root microbiota on fungal and oomycetal community structure and diversity. We demonstrate that the bacterial microbiota is essential for plant survival and protection against root-derived filamentous eukaryotes. Deconvolution of 2,862 binary bacterial-fungal interactions *ex situ*, combined with community perturbation experiments *in planta*, indicate that biocontrol activity of bacterial root commensals is a redundant trait that maintains microbial interkingdom balance for plant health.

## Introduction

Similar to the guts of vertebrates, the roots of soil-grown plants are inhabited by taxonomically structured bacterial communities that provide fitness benefits to their respective hosts (Hacquard et al., 2015). A possibly unique feature of plant roots is their capacity to host simultaneously, besides the bacterial microbiota (Lebeis et al., 2015; Bai et al., 2015; Castrillo et al., 2017), also a range of soil-borne filamentous eukaryotic microbes such as fungi and oomycetes that have evolved independently in distinct kingdoms of life (Mycota and Chromista; Ruggiero et al., 2015). The dynamics of microbe-microbe interactions have recently emerged as an important feature of the phyllosphere (Agler et al., 2016), and such interactions are now acknowledged to carry out important functions for plant health, including synergistic effects on plant growth (van der Heijden et al., 2016), protection against microbial pathogens (Santhanam et al., 2015), and promotion of mycorrhizal symbiosis (Garbaye et al., 1994). In contrast, the interplay between prokaryotic and eukaryotic microbes along the soil-root continuum and the relevance of inter-kingdom microbe-microbe interactions for structuring root-associated microbial communities have received little attention so far (Hassani et al., 2018).

By profiling three microbial groups (bacteria, fungi, oomycetes) in the roots of natural *A. thaliana* populations and establishing reference microbial culture collections for microbiota reconstitution experiments, we provide community-level evidence that negative interactions between prokaryotic and eukaryotic root microbiota members are critical for plant host survival and maintenance of host-microbiota balance.

## Results

### Root-associated microbial assemblages

We collected *A. thaliana* plants from natural populations at two neighbouring sites in Germany (Geyen and Pulheim; 5 km apart) and a more distant location in France (Saint-Dié; ~300 km away) (**Figure S1**; **Table S1**). For each population, four replicates, each consisting of four pooled *A. thaliana* individuals were prepared, together with corresponding bulk soils. Root samples were fractionated into episphere and endosphere compartments, enriching for microbes residing on the root surface or inside roots, respectively (**Figure S2**). We characterized the multi-kingdom microbial consortia along the soil-root continuum by simultaneous DNA amplicon sequencing of the bacterial 16S rRNA gene and fungal as well as oomycetal Internal Transcribed Spacer (ITS) regions (Agler et al. 2016) (**Table S2**). Alpha diversity indices (within-sample diversity) indicated a gradual decrease of microbial diversity from bulk soil to the root endosphere (Kruskal-Wallis test, *p*<0.01; **Figure S3**). Profiles of microbial class abundance between sample-types (**Figure 1A**) and Operational Taxonomic Unit (OTU) enrichment tests conducted using a linear model between soil, root episphere and root endosphere samples (*p*<0.05, **Figure 1B**) identified 96 bacterial, 24 fungal and one oomycetal OTU that are consistently enriched in plant roots across all three sites. This, together with the reduced alpha diversity, points to a gating role of the root surface for entry into the root interior for each of the three microbial kingdoms (Bulgarelli et al., 2012; Lundberg et al., 2012; Edwards et al., 2015). The root-enriched OTUs belong to the bacterial classes Gammaproteobacteria (34%), Actinobacteria (26%), Betaproteobacteria (24%), the fungal classes Dothideomycetes (40%) and Sordariomycetes (21%) and the oomycetal genus Pythium (**Figure 1B**). By examining between-sample variation (beta-diversity, Bray-Curtis distances), we found that the bacterial communities cluster along the first PCoA axis according to the compartment, whereas the major factor explaining fungal and oomycetal communities is host biogeography (**Figure 1C**). Microbial co-occurrence network analysis (**Figure S4**) and permutational multivariate analysis of variance (PERMANOVA test, see methods) corroborated the contrasting effects of compartment and site on the structure of bacterial (compartment: 41%, *p*<0.001; site: 19%, *P* < 0.001), fungal (compartment: 21%, *p*<0.001; site: 43%, *p*<0.001), and oomycetal (compartment: 14% *P* <0.001; site: 37% *P* <0.004) communities. Although the different rates of divergence of the marker loci used to characterize community composition of the tested microbial kingdoms might inflate these striking differences between root-associated prokaryotic and eukaryotic microbial communities, our results suggest that geographically distant *A. thaliana* populations host taxonomically similar root-associated bacterial communities, but more dissimilar and site-specific root-associated fungal and oomycetal assemblages (Coleman-Derr et al., 2016).

**Figure 1:**
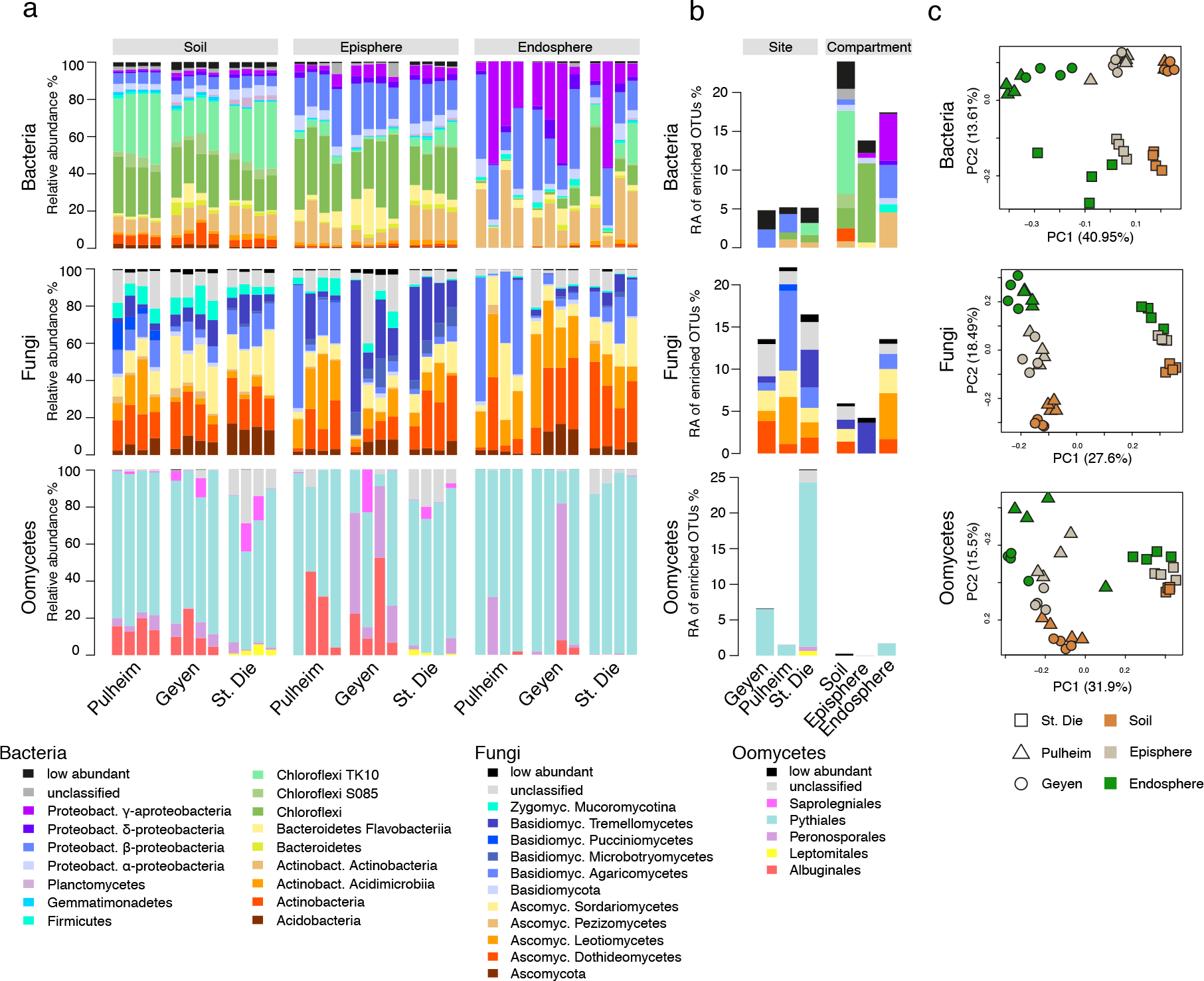
Microbial community structure in three natural *A. thaliana* populations. **a,** Relative abundance of bacterial, fungal and oomycetal taxa in soil, root episphere and root endosphere compartments in three sites (Pulheim and Geyen in Germany and Saint-Dié in France). The taxonomic assignment is based on the RDP using a bootstrap cutoff of 0.5%. Low- abundance taxonomic groups with less than 0.5% of total reads across all samples are highlighted in black. Each technical replicate comprised a pool of four plants. **b,** Relative abundance (RA) of OTUs significantly enriched in a specific site or compartment. A generalized linear model was used to compare OTU abundance profiles in one site or compartment versus the other two sites or compartments, respectively (*p* < 0.05, FDR corrected). The relative abundances for these OTUs were aggregated at the class level. **c,** Community structure of bacteria, fungi and oomycetes in the 36 samples was determined using principal component analysis. The first two dimensions of a principal component analysis are plotted based on Bray-Curtis distances. Samples are colour-coded according to the compartment and sites are depicted with different symbols.

### Root microbiota cross-kingdom connectivity

To investigate intra- and inter-kingdom microbial OTU relationships, we used the network inference tool SparCC (Friedman and Alm, 2012) and performed compartment-specific network analysis using community profiling data from the aforementioned three natural sites (**Figure 2; Figures S5 and S6**). To reduce the influence of site-specific OTUs on network structure, we selected core microbial OTUs shared in >80% of either bulk soil, episphere or endosphere samples across all three sites with relative abundances of >0.1%. Consistent with alpha diversity indices (**Figure S3**), microbial network complexity decreases from soil to endosphere compartments (soil/episphere/endosphere: 731/454/178 OTUs and 42,043/8,518/1,125 edges, respectively; **Figure 2; Figures S5 and S6**). Notably, inspection of root network architecture indicates that correlations between bacterial and fungal OTUs are primarily negative (92.0%), whereas positive correlations dominate within bacterial (87.7%) and fungal (89.7%) groups (**Figure 2A** and **2B**). The most positively and negatively correlated OTUs defined with the SparCC method were independently validated using Spearman correlation (**Figure S7**). To test for potential phylogenetic signal(s) in the correlation network, we calculated the strength of bacterial-fungal correlations for taxonomic groups comprising at least 5 OTUs (**Figure 2C**). This revealed that bacterial OTUs belonging to the classes Betaproteobacteria, Bacilli and Deltaproteobacteria, as well as fungal OTUs belonging to the classes Dothideomycetes, Leotiomycetes and Tremellomycetes display the strongest negative correlations with fungal and bacterial OTUs, respectively. Further inspection of fungal and bacterial OTUs that display the highest betweenness centrality scores and the highest negative cross-kingdom connection frequencies identified 5 OTUs belonging to the fungal genera Davidiella and Alternaria and the bacterial genera Variovorax, Kineosporia and Acidovorax that might represent keystone taxa driving fungal-bacterial balance in *A. thaliana* roots (**Figure 2D**). Collectively, our network analysis indicates strong negative correlations between several bacterial and fungal taxa in plant roots.

**Figure 2:**
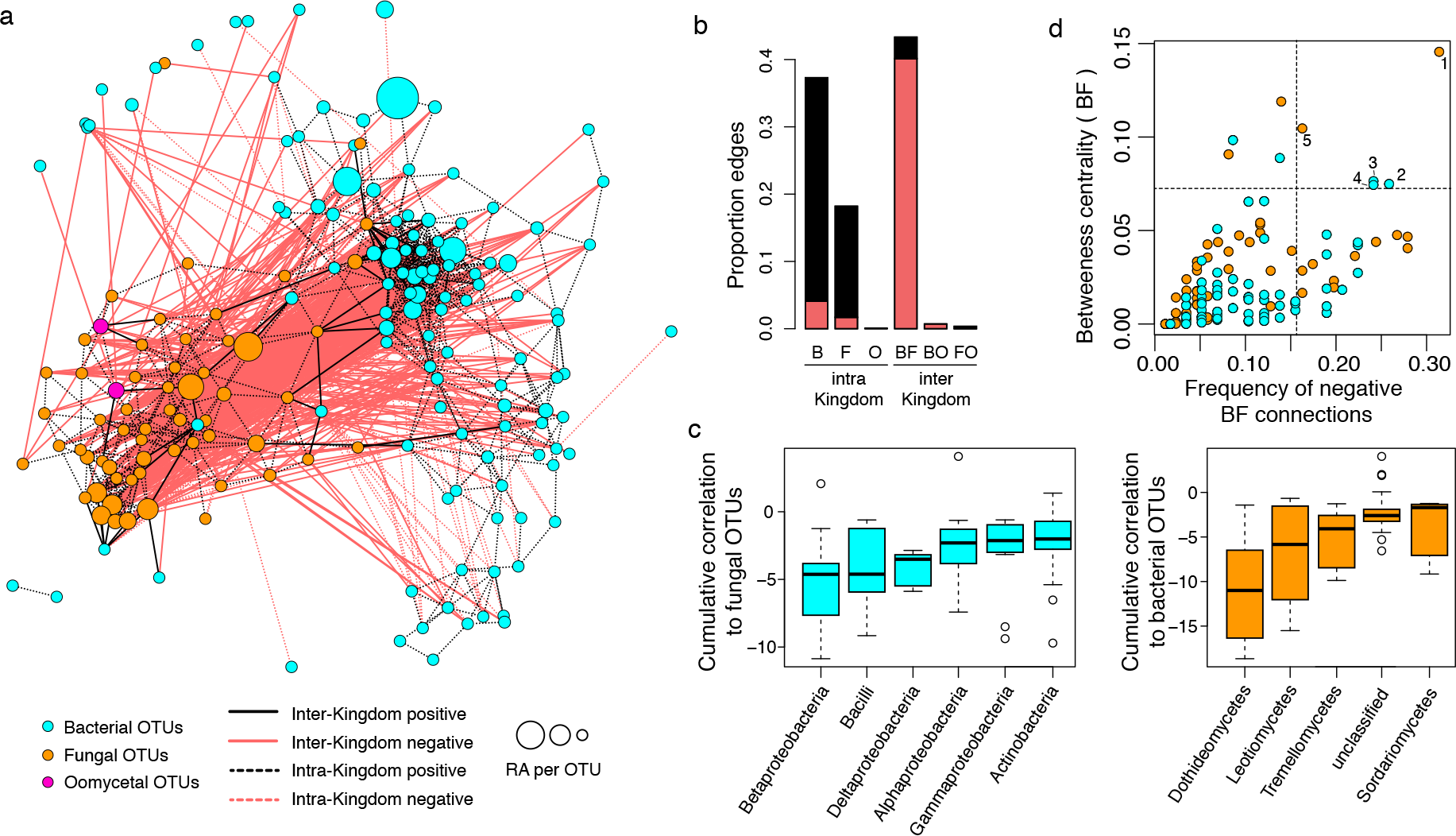
Microbial network of the *A. thaliana* root endosphere microbiota. **a,** Correlation-based network of root-associated microbial OTUs detected in three natural *A. thaliana* populations (Pulheim, Geyen, Saint-Dié). Each node corresponds to an OTU and edges between nodes correspond to either positive (black) or negative (red) correlations inferred from OTU abundance profiles using the SparCC method (pseudo *p*-value >0.05, correlation values <-0.6 or >0.6). OTUs belonging to different microbial kingdoms have distinct colour codes and node size reflects their relative abundance (RA) in the root endosphere compartment. Intra-kingdom correlations are represented with dotted lines and inter-kingdom correlations by solid lines. **b,** Proportion of edges showing positive (black) or negative (red) correlations in the microbial root endosphere network. B: bacteria, F: fungi, O: oomycetes. **c,** Cumulative correlation scores measured in the microbial network between bacterial and fungal OTUs. Bacterial (left) and fungal (right) OTUs were grouped at the class level (> five OTUs/class) and sorted according to their cumulative correlation scores with fungal and bacterial OTUs, respectively. **d,** Hub properties of negatively correlated bacterial and fungal OTUs. For each fungal and bacterial OTU, the frequency of negative interkingdom connections is plotted against the betweenness centrality inferred from all negative BF connections (cases in which a node lies on the shortest path between all pairs of other nodes). The five microbial OTUs that show a high frequency of negative inter-kingdom connections and betweenness centrality scores represent hubs of the “antagonistic” network and are highlighted with numbers. 1: Davidiella; 2: Variovorax; 3: Kineosporia; 4: Acidovorax; 5: Alternariab

### Root-derived microbial culture collections

To test whether cross-kingdom microbe microbe competition is relevant along the soil-root continuum, we first deconstructed the microbiota of *A. thaliana* roots by generating microbial culture collections of root-associated bacteria, fungi and oomycetes. For bacteria, we used the recently established *A. thaliana* root-derived bacterial culture collection, in which 54-65% of bacterial root-enriched OTUs have one or several isolates in pure culture (Bai et al., 2015). To complement this collection, we conducted a large-scale isolation of filamentous eukaryotic microbes from the roots of 24 *A. thaliana* Col-0 individuals at three developmental stages and *A. thaliana* relatives grown in Cologne Agricultural Soil (CAS), the same soil that was used to establish the root-derived bacterial culture collection^3^. Additionally, two to six plant individuals, grown in the Geyen, Pulheim and Saint-Dié natural sites, were used for the isolation of root-associated fungi and oomycetes. In total, ~12,000 surface-sterilized root segments (~5 mm each) were placed on five agar-based growth media, resulting in the purification of 202 filamentous eukaryotes (**Table S3**). Taxonomic assignment based on Sanger sequencing of the fungal or oomycetal ITS was successful for 176 isolates, of which 93% are fungi and 7% oomycetes (**Table S3**). After elimination of potential clonal duplicates, defined as microbes with 100% identical ITS retrieved from plant roots grown in the same soil, the fungal culture collection comprises 69 isolates that belong mainly to the classes Sordariomycetes (65%), Dothideomycetes (24%) and Agaricomycetes (4%), whereas the oomycete collection contains 11 isolates that exclusively belong to the genus Pythium (**Figure S8**). The results from our culture-dependent approach mirror the taxonomic composition of root-associated fungal and oomycetal communities from natural *A. thaliana* populations determined using culture-independent ITS amplicon sequencing (**Figure 3A**). To estimate recovery rates, we compared the Sanger-based reference ITS sequences with the corresponding culture-independent datasets (> 0.1% RA) at 97% sequence similarity (**Figure 3B**). By considering abundant OTUs that together represent 60% or 80% of the total number of sequence reads detected in all root samples, we estimate recovery rates of 50% and 37% for fungi and 50% and 28% for oomycetes, respectively (**Figure 3B**). It is likely that some root-derived filamentous eukaryotes are recalcitrant to isolation because of their obligate biotrophic lifestyle such as *Olpidium brassicae* corresponding to the abundant OTU 8 (**Figure 3B**). Our findings show that several of the most abundant filamentous eukaryotes associated with the roots of *A. thaliana* plants grown in natural populations can be retrieved as pure cultures, providing opportunities for a holistic reconstitution of the root microbiota under laboratory conditions.

**Figure 3:**
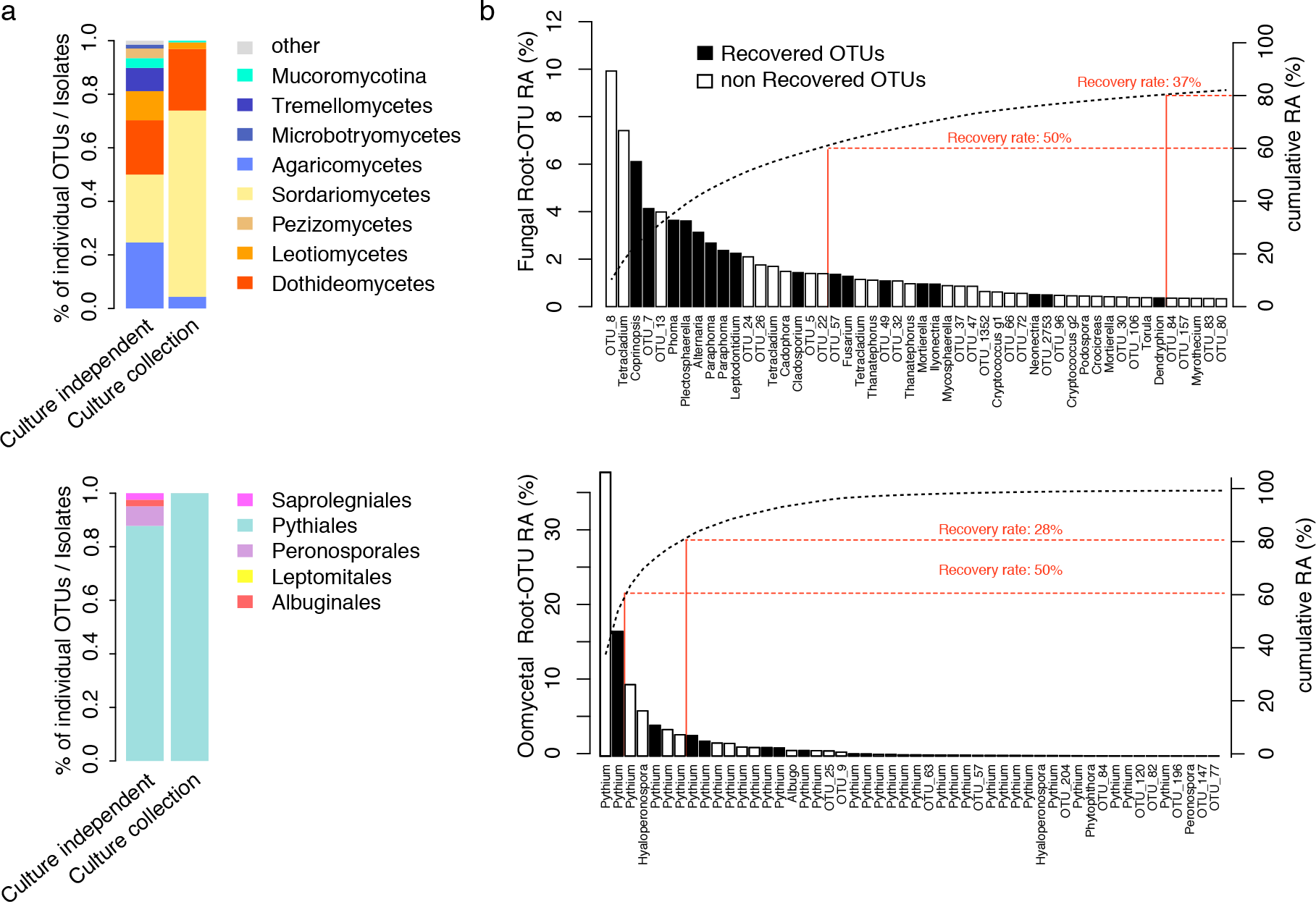
Recovery rates and taxonomic representation of root-derived fungal and oomycetal culture collections. **a,** Comparison of fungal (upper panel) and oomycetal (lower panel) taxonomic composition between culture-dependent and culture-independent methods. Culture collection: taxonomic composition (class level) of the 69 fungal and 11 oomycetal strains isolated from plant roots grown in the Cologne agricultural soil and the three natural sites Pulheim, Geyen, and Saint-Dié (**Figure S8**). Culture-independent approach: taxonomic composition of fungal and oomycetal root-associated OTUs (> 0.1% RA in at least one site, RDP bootstrap at class level >=0.8) detected in the roots of *A. thaliana* grown in the same soils used for the culture-dependent approach. b, Recovery rates of fungal (upper panel) and oomycetal (lower panel) isolates from the culture collections at different thresholds. The rank abundance plots show the 50 most abundant root-associated fungal and oomycetal OTUs from *A. thaliana* grown in the above-mentioned soil types, together with their cumulative relative abundance (RA). OTUs that have a representative isolate in the culture collections (97% sequence similarity) are highlighted with black bars. The percentages of naturally occurring OTUs recovered as pure cultures are given for OTUs representing 60% and 80% of the total read counts.

### Multi-kingdom root microbiota reconstitution

Using microbes that were exclusively recovered from the roots of *A. thaliana* or close relatives grown in CAS soil, we assembled seven complex synthetic microbial communities consisting of 148 bacteria (B), 34 fungi (F) or nine oomycetes (O) as well as all corresponding combinations of multi-kingdom microbial consortia (BO, BF, FO, BFO; **Table S4**). These microbes, selected based on their taxonomic diversity, were used to re-populate the gnotobiotic FlowPot system containing peat and vermiculite as sterile soil matrix onto which surface-sterilized *A. thaliana* Col-0 seeds were placed (Kremer et al., 2018). Upon co-incubation of these microbial communities and the plant host for four weeks in this substrate, the filamentous eukaryotic microbes in the absence of bacterial root commensals (F, O, FO) had a strongly detrimental impact on plant growth (p<0.05, Kruskal-Wallis with Dunn’s post-hoc tests) and their survival rate, whereas in combination with the bacterial community (BO, BF), plant growth was rescued to similar levels as in microbe-free (MF) control plants (**Figure 4A**). Interestingly, increasing the complexity of the microbial consortium (BFO) resulted in significant plant growth promotion (125% of plant biomass compared to MF plants; p<0.05, Kruskal-Wallis with Dunn’s post-hoc tests), indicating beneficial activities of multi-microbial consortia for plant growth (**Figure 4A**). To examine whether microbial interactions contribute to the observed plant fitness phenotypes, we simultaneously profiled bacterial, fungal and oomycetal communities established in the roots and the surrounding FlowPot matrix using the same sequencing strategy described above for natural site samples. Consistent with the hypothesis that bacteria restrict fungal and oomycetal growth along the soil-root continuum, the presence of the bacterial community significantly reduced the alpha diversity of these groups in the matrix (Kruskal-Wallis with Dunn’s post-hoc tests, p<0.05, **Figure 4B**; **Figure S9**). Remarkably, the taxonomic structure of the bacterial communities remained essentially unaltered in both matrix and root compartments in the presence of filamentous eukaryotic microbes (B vs. BF, BO, BFO), whereas clear fungal and oomycetal community shifts were observed in the presence of bacteria (F, FO vs. BF, BFO / O, FO vs. BO, BFO; **Figure 4C**; **Figure S10**). Importantly, the bacteria-mediated fungal community shift is largely plant-independent as this shift was also seen in the unplanted matrix (BFO UNPL). PERMANOVA indicates that 11.6% and 7.8% of the variance in the fungal and oomycetal community structure, respectively, is explained by the presence of bacteria, whereas the presence of filamentous eukaryotes explains only 2.20% to 3.65% of the bacterial community structure (p<0.05, **Table S5**). Enrichment tests conducted using a linear model identified 11 fungal and four oomycetal isolates for which the relative abundance is significantly increased in the absence of bacteria in matrix samples (**Figure S11**). Given that the bacteria-mediated fungal and oomycetal community shifts are linked to plant growth rescue, this indicates that a major physiological function of the bacterial root microbiota is to protect plants from the detrimental activities of root-derived filamentous eukaryotes.

**Figure 4:**
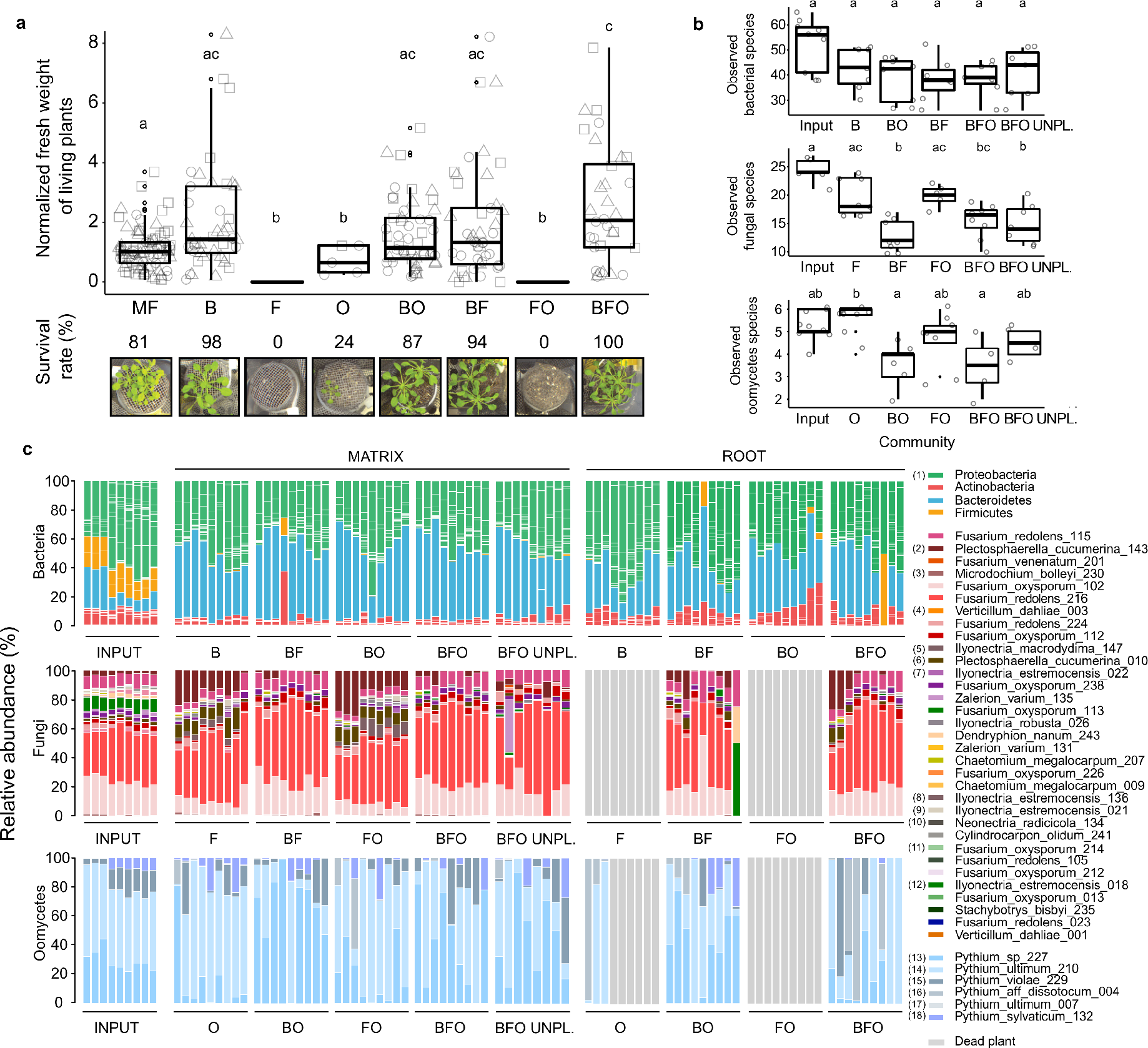
Multi-kingdom reconstitution of the *A. thaliana* root microbiota. **a,** Recolonization of germ-free plants with root-derived bacterial (148), fungal (34) and oomycetal (9) isolates in the FlowPot system. Shoot fresh weight of four-week-old *A. thaliana* Col-0 inoculated with bacteria (B), fungi (F), oomycetes (O) bacteria and oomycetes (BO), bacteria and fungi (BF), fungi and oomycetes (FO) and bacteria, fungi and oomycetes (BFO). MF: microbe-free/control. Shoot fresh weight values were normalized to MF. Significant differences are depicted with letters (p<0.05, Kruskal-Wallis and Dunn’s *post-hoc* tests). Survival rate values represent the percentage of germinated plants that survived. Data are from three biological replicates (represented by different shapes) with three technical replicates each. **b,** Observed species per microbial group in matrix samples for each of the above-mentioned inoculations (*p*<0.05, Kruskal Wallis and Dunn’s *post-hoc* tests). Input: initial microbial inoculum. UNPL: unplanted matrix. **c,** Relative abundances of microbial isolates in initial input and output matrix and root samples after four weeks. Taxonomic assignment is shown at the phylum level for bacteria and at the species level for fungi and oomycetes. Numbers in brackets refer to enriched species in **Figure S11B**.

### Redundancy in bacterial biocontrol activity

To clarify whether the detrimental effect on plant growth observed with the 34-member fungal community is mediated by one or several fungal isolates, we performed re-colonization experiments with individual fungal isolates with the gnotobiotic FlowPot system (**Figure S12**). This revealed that a majority, i.e. 18/34, of root-derived fungi isolated from healthy *A. thaliana* restrict plant growth in mono-associations with the host, whereas non-significant effects on plant growth were found for 16/34 fungal strains (*p*<0.01, Kruskal-Wallis, Dunn’s post-hoc test). The 11 fungal isolates that show significantly higher relative abundance in the absence of the bacterial root microbiota (see above) include both pathogenic and non-pathogenic fungi and a 23-member fungal community lacking these 11 isolates remains harmful for plant growth (**Figure S12**).

To identify bacterial taxa potentially contributing to plant growth rescue in our multi-kingdom microbiota reconstitution experiment, we developed a high-throughput *ex situ* bacterial-fungal interaction screen (**Figure 5A**; **Figure S13**). In brief, spores collected from sporulating fungal isolates were distributed into 96-well plates containing liquid TSB medium (20%) with or without individual bacterial root commensals for 48 hours. Fungal growth was determined by fluorescence using a chitin binding assay (Figueroa-López et al., 2014; **Figure S13**). We tested 2,862 binary interactions in triplicate with the aforementioned individual bacterial root commensals against a phylogenetically diverse set of 27 fungi from our culture collection, including seven that were included in the microbiota reconstitution experiment (**Figure 5A**; **Table S6**). We identified clear phylogenetic signals among the detected antagonistic interactions, including several bacterial strains belonging to the families Comamonadaceae, Pseudomonadaceae, Rhizobiaceae, and Flavobacteriaceae that exhibit a high competitive potential. Therefore, several taxonomic lineages of the bacterial root microbiota can exert direct inhibitory activities against a wide range of *A. thaliana* root-associated fungi. Most actinobacterial strains show comparatively weak or insignificant inhibitory activity against the tested fungal isolates. Comparison of correlation strength, defined by SparCC network analysis with samples from the natural sites (**Figure 2**), with the binary interaction phenotypes determined in the high-throughput fungal-bacterial interaction screen revealed consistent trends for most tested bacterial lineages and identified Variovorax and Acidovorax strains lineage (Comamonadaceae family) as promising biocontrol microbes (**Figure S14**). This suggests that direct antifungal activities of bacterial root commensals are important determinants underlying the largely negative correlations between bacterial and fungal OTUs in root-associated multi-kingdom microbial communities of plants grown in nature.

**Figure 5:**
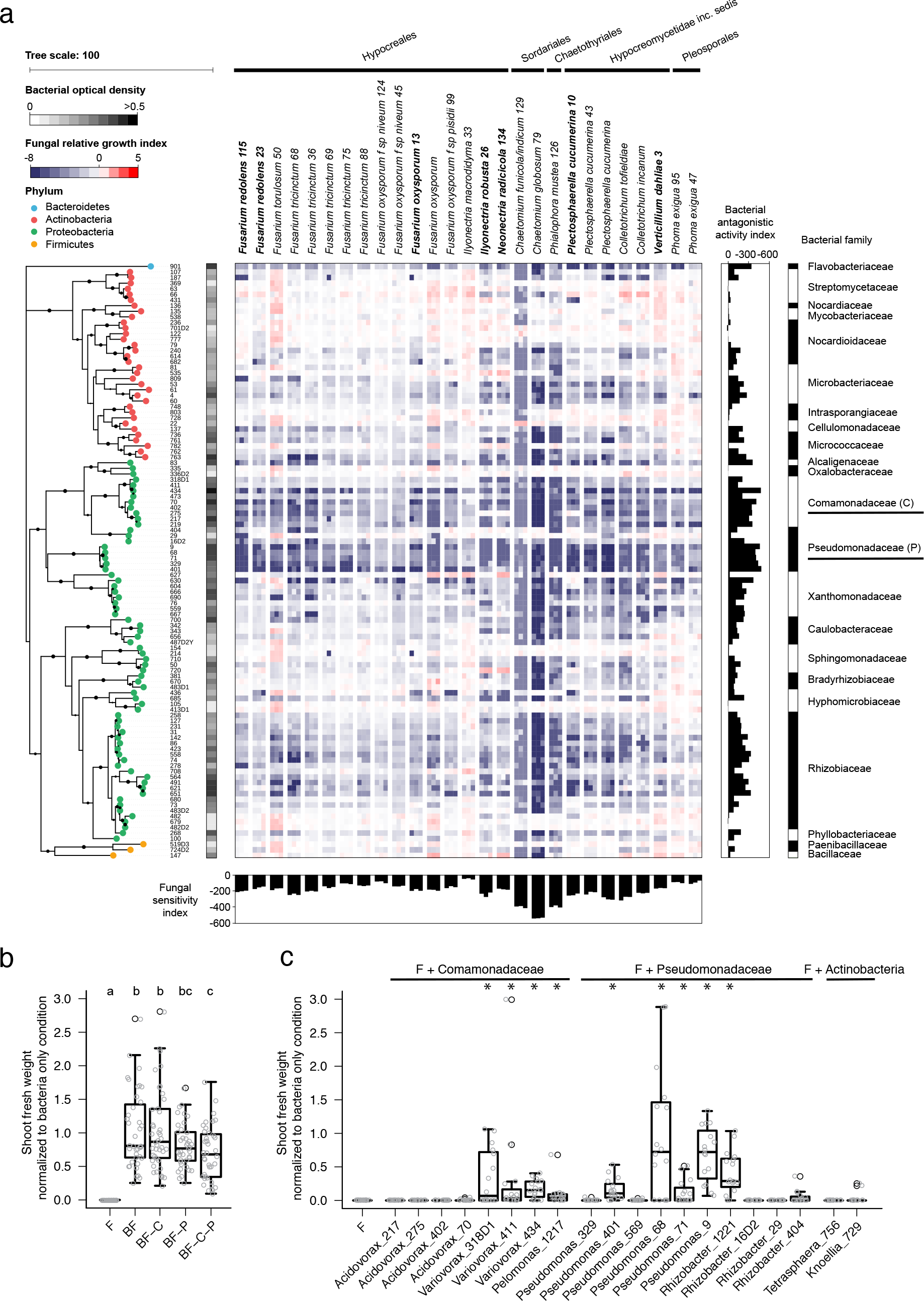
Inhibitory activities of bacterial root microbiota members towards root-associated fungi. **a,** Alteration of fungal growth upon interaction with phylogenetically diverse bacterial root commensals. The heatmap depicts the log2 fungal relative growth index (presence vs. absence of bacterial competitors) measured by fluorescence using a chitin binding assay against Alexa Fluor 488-conjugated WGA. The phylogenetic tree was constructed based on the full bacterial 16S rRNA gene sequences and bootstrap values are depicted with black circles. Vertical and horizontal barplots indicate the cumulative antagonistic activity for each bacterial strain and the cumulative sensitivity score for each fungal isolate, respectively. Alternating white and black colours are used to distinguish the bacterial families. All bacteria presented and 7/27 fungi (highlighted in bold) were used for the above-mentioned multi-kingdom reconstitution experiment (see **Figure 4**). **b,** Shoot fresh weight of fungi and bacteria-inoculated plants relative to the bacteria-only inoculated plants in depletion experiments, in which specific bacterial families (C: *Commonadaceae*; P: *Pseudomonadaceae*w) were removed to test their fungal control capacity. Significant differences are depicted with letters (Kruskal-Wallis with Dunn’s *post-hoc* test, *p*<0.05). **c,** Same experiment as in b, but instead of depleting strains, single bacterial isolates were coinoculated with the 34-member fungal community (F) to test their plant growth rescue activities.

We reasoned that if bacteria-mediated fungal growth inhibition detected in our binary screen is maintained in a community context, then depletion of the most competitive bacterial isolates belonging to the families Pseudomonadaceae (-P, 8 members) and/or Comamonadaceae (-C, 10 members) from the full 148-member bacterial community might result in a reduction of bacterial anti-fungal activity. We re-colonized germ-free *A. thaliana* with the aforementioned 34-member fungal community (F) in the presence of the full or three perturbed bacterial consortia (BF or BF-C, BF-P, BF-C-P) in the FlowPot system (**Figure 5B**). Community profiling of bacterial outputs validated the withdrawal of all Pseudomondaceae isolates in the −P and −CP samples and depletion of 9 out of 10 Comamonadaceae isolates in the −C and −CP samples, leading to a 59% reduction of Comamonadaceae-related reads in depleted vs. non-depleted samples (**Figures S15 and S16**). The bacterial community lacking bacterial strains from these two families, but not those depleted of either Pseudomonadaceae or Comamonadaceae family members, failed to fully rescue plant growth to control levels in the presence of the synthetic fungal community (BF vs BF-C-P, Kruskall Wallis with Dunn’s post-hoc tests, *p*<0.05, **Figure 5B**), suggesting that bacterial root commensals of the families Pseudomonadaceae and Comamonadaceae contribute to the observed plant growth rescue. Notably, several individual bacterial strains from these two families, belonging to the genera Variovorax (3/3), Pseudomonas (4/6), Pelomonas (1/1), and Rhizobacter (1/4), but not Acidovorax (0/4), were sufficient to fully or partially rescue fungus-mediated plant growth inhibition in the presence of the 34-member fungal community (**Figure 5C**). In conclusion, network-based predictions of root microbiota cross-kingdom connectivity with samples collected from natural sites, binary fungal-bacterial interaction assays and community perturbation experiments with germ-free plants, indicate that the protective activity conferred by the bacterial root microbiota is a redundant trait mediated by several taxonomic lineages of the bacterial root microbiota. This redundancy is likely important in conferring robust host protection and maintaining host-microbiota homeostasis.

## Discussion

Bacteria, fungi and oomycetes arose ~3,500, ~1,050 and ~500 million years ago (MyA), respectively, and likely co-existed and interacted in soils before plants colonized terrestrial habitats ~450 MyA (Hassani et al., 2018). The likely competition between these microbial groups for soil organic matter, including root-secreted photoassimilates (Zhalnina et al. 2018) could explain our finding that positive correlations dominate within a microbial kingdom, whilst negative correlations dominate between bacteria and the filamentous eukaryotes.

Although our root-derived microbial culture collections still lack some abundant microbiota members, the composition of CAS-derived synthetic bacterial and fungal communities established in the roots of germ-free *A. thaliana* plants more closely resemble those of plant roots grown in the corresponding CAS soil than in the other tested natural soils (**Figure S17**). This suggests that the pronounced impact of root-associated bacteria on fungal and oomycetal community structure in our gnotobiotic plant system (explaining >7 and >10% of microbial interkingdom variance, respectively) partially recapitulates microbial interactions in the natural environment that are necessary for plant survival. Given that *A. thaliana* root-associated fungal and oomycetal communities display strong biogeographical signatures, likely influenced by their dispersal limitation and/or climate (Coleman-Derr et al., 2016; Talbot et al., 2014), we propose that biocontrol activities of bacterial root commensals towards phylogenetically unrelated fungi is relevant to confer robust plant protection. The bacterial isolates belonging to the genus Variovorax and Pseudomonas that confer robust host protection are members of the core root microbiota (Hacquard et al., 2015) and represent 24% of the 16S rRNA reads detected in *A. thaliana* roots across the three natural sites. Our observation that individual bacterial strains, belonging to distinct taxonomic lineages of the bacterial root microbiota, are sufficient to protect the plant host against taxonomically diverse root-colonizing fungi provides a rational framework to explain at least part of the activity of biocontrol bacteria used in field applications in agricultural contexts. The lack of comprehensive microbial culture collections from unplanted CAS soil did not allow us to directly test whether the bacterial root microbiota, which is horizontally acquired from a small fraction of the bacterial soil biome (Bulgarelli et al., 2012; Lundberg et al., 2012), is enriched for members that restrict root colonization by filamentous eukaryotes. However, this hypothesis is indirectly supported by our network-based microbial interkingdom analysis, in which the ratio of negative to positive correlations between prokaryotic and filamentous eukaryotic microbes shifted from 4:1 in the soil network to 12:1 in the root network (**Figures 2, S5, and S6**). Hence, we conclude that the detected microbial interkingdom interactions take place at the soil-root interface during microbiota establishment and are maintained inside plant roots, which might explain the similarities in microbial interkingdom community shifts in roots and unplanted peat matrix samples in the microbiota reconstitution experiments with microbes derived exclusively from roots. Although these similarities potentially imply that plant host-derived cues are dispensable for the antagonistic activity of the bacterial root microbiota against filamentous eukaryotic microbes, initial establishment of the bacterial root microbiota by the plant host might still be necessary.

Given that all bacterial, fungal, and oomycetal strains used in our study were isolated from roots of healthy *A. thaliana* plants, the contrasting effects of synthetic communities comprising bacteria and filamentous eukaryotes on plant health are surprising. Loss of mycorrhiza symbiosis in *A. thaliana* or relatives appears to have been partly compensated by associations with other beneficial fungal root endophytes (Almario et al., 2017; Hiruma et al., 2016; Hacquard et al., 2016). Our data show that roots of *A. thaliana* in their natural habitats host a rich diversity of filamentous eukaryotes, dominated by Ascomycetes, but also demonstrate that in the absence of bacterial competitors, consortia of filamentous root-derived eukaryotes (F, O, FO) have chiefly detrimental activities on plant health and survival. Strikingly, >50% of the isolates restrict plant growth in mono-associations with the plant host. This is consistent with earlier reports (Keim et al., 2014; Kia et al., 2017) and suggests that numerous *A. thaliana* root-associated fungi and oomycetes cannot be kept at bay by the plant innate immune system alone. However, re-colonization of *A. thaliana* with the most complex multi-kingdom microbial consortium (BFO) resulted in maximal plant growth and survival in our gnotobiotic plant system. Thus, we propose that mutual selective pressures, acting on the plant host and its associated microbial assemblage, have, over evolutionary time scales, favoured interkingdom microbe-microbe interactions rather than associations with a single microbial class.

## Experimental procedures

### Sampling of *A. thaliana* plants in their natural habitats

*A. thaliana* plants were harvested from three natural populations: two in Germany (Geyen, Pulheim) and one in France (Saint-Dié). For each population, 16 plant individuals were dug out with their surrounding soil, transferred into sterile falcons, and transported on ice to the laboratory. Sample fractionation into soil, root episphere and root endosphere compartments was performed within 12 hours after harvesting (**Figure S2**). Soil particles not in contact with roots were transferred to 2-ml Lysis Matrix E tubes (MP Biomedicals, Solon, USA) and are defined as the soil fraction. Plant roots were cut and thoroughly washed with sterile water to remove visible soil particles. Epiphytic microbes were washed away from root systems using extensive shaking in TE buffer supplemented with 0.1% Triton. These washes were filtered through 0.22-μM pore size membranes and considered as epiphytic fraction. Root systems were then washed successively in 80% EtOH and 0.25% NaOCl to further clean the root surfaces from living microorganisms, and subsequently washed three times (1 min each) in sterile water. These microbially-enriched endosphere fractions were transferred to 2-ml tubes. Each of the four biological replicates consists of a pool of four plant individuals.

### Microbial community profiling from natural sites

Total DNA was extracted from aforementioned samples, using the FastDNA® SPIN Kit for Soil (MP Biomedicals, Solon, USA). Samples were homogenized in Lysis Matrix E tubes using the Precellys®24 tissue lyzer (Bertin Technologies, Montigny-le-Bretonneux, France) at 6,200 rps for 30 seconds. DNA samples were eluted in 60 μl nuclease-free water and used for bacterial, fungal and oomycetal community profiling (Agler et al. 2016). Concentration of DNA samples was fluorescently quantified, diluted to 3.5ng/uL, and used in a two-step PCR amplification protocol. In the first step, V4-V7 of bacterial 16s rRNA (799F - 1192R), fungal ITS1 (ITS1F - ITS2) and oomycetal ITS1 (ITS1-O - 5.8s-Rev-O) (**Table S2**) were amplified. Under a sterile hood, each sample was amplified in triplicate in 25 μl reaction volume containing 2 U DFS-Taq DNA polymerase, 1x incomplete buffer (both Bioron GmbH, Ludwigshafen, Germany), 2mM MgCl2, 0.3% BSA, 0.2 mM dNTPs (Life technologies GmbH, Darmstadt, Germany) and 0.3 μM forward and reverse primer PCR was performed using the same parameters for all primer pairs (94 °C/2 minutes, 94 °C/30 seconds, 55 °C/30 seconds, 72 °C/30 seconds, 72 °C/10 minutes for 25 cycles). Afterwards, single-stranded DNA and proteins were digested by adding 1 μl of Antarctic phosphatase, 1 μl Exonuclease I and 2.44 μl Antarctic phosphatase buffer (New England BioLabs GmbH, Frankfurt, Germany) to 20 μl of the pooled PCR product. Samples were incubated at 37°C for 30 minutes and enzymatic activity was deactivated at 85°C for 15 minutes. Samples were centrifuged for 10 minutes at 4,000 rpm and 3 μl of this reaction was used for a second PCR, where cycles were reduced to 10 and with primers including barcodes and Illumina adaptors (**Table S2**). PCR reactions were prepared in the same way as described above using the same protocol except the number of PCR-cycles were reduced to 10. PCR quality was controlled by loading 5 μl of each reaction through a 1% agarose gel and affirming that no band within the negative control was detected. Afterwards, the replicated reactions were combined and purified: 1) bacterial amplicons were loaded in a 1.5% agarose gel and ran for 2 hours at 80 V; bands with the correct size of ~500 bp were cut and purified using the QIAquick gel extraction kit (Qiagen, Hilden, Germany); 2) fungal and oomycetal amplicons were purified using Agencourt AMPure XP beads. DNA concentration was again fluorescently determined, and 30 ng DNA of each of the barcoded amplicons were pooled in one library per microbial group. Each library was then purified and re-concentrated twice with Agencourt AMPure XP beads and 100ng of each library was pooled together. Paired-end Illumina sequencing was performed in-house using the MiSeq sequencer and custom sequencing primers (**Table S2**).

### 16S and ITS read processing

Paired 16s rRNA amplicon sequencing reads were joined (join_paired_ends Qiime, default) and then quality-filtered and demultiplexed (split_libraries_fastq, Qiime, with max. barcode errors 1 and phred score of 30; Caporaso et al., 2010). The filtered reads were dereplicated (usearch-derep_fulllength), sorted by copy number (only reads >2 copies were retained) and clustered using the usearch algorithm at 97% sequence identity (Edgar, 2013). Clustered reads were checked for chimeras using usearch (usearch -uchime_ref, gold db ?!). All retained OTUs were aligned to the greengenes db (DeSantis et al., 2006) using PyNAST (Caporaso et al., 2009); those that did not align were removed. To each OTU a taxonomic classification was assigned using Qiime (assign_taxonomy from qiime, uclust algorithm with default param, greengenes db). Mitochondria-assigned OTUs were eliminated. Out of the remaining sequences an OTU table was built (usearch_global 97%,). ITS amplicon data were processed as followed. Reads were joined and demultiplexed as described in the previous section, and forward reads were also demultiplexed and filtered. For reads where no joined pair of reads exists the forward reads were kept. The combined reads were trimmed to an length of 220bp. Reads were dereplicated and sorted (keeping only those with >2 copies). The presence of ITS sequences was determined using ITSx (Bengtsson-Palme et al., 2013) and reads containing ITS sequences were clustered at 97% using usearch. Fungal OTUs were checked for chimeric sequences using (uchime_ref) against a dedicated chimera detection db (Nilson et al., 2015). Oomycetal OTUs were checked using the -chime_denovo function from usearch. To check for non-fungal/non-oomycetal sequences, the remaining OTU sequences were blasted against an ITS-sequence database. For this purpose, all available ITS sequences (search term “internal+transcribed”, for plants, animals, fungi, oomycetes and protitst) were retrieved from the NCBI nucleotide database (January/February 2016). All OTU sequences whose best blast hit (bbh) was not annotated as a fungal/oomycetal sequence were removed, as were all sequences that showed more hits in non-fungal/non-oomycetal sequences (out of max. 10 hits). Taxonomic classification was done with the rdp classifier (Wang et al., 2007) using the warcup database for fungal OTUs (Deshpande et al., 2016) and a self-established db for oomycetal OTUs. The latter was constructed from NCBI-derived ITS sequences (checked using ITSx, remaining sequences were used to train the RDP classifier).

### Calcul of alpha- and beta-diversity indices

To assess alpha-diversity within natural samples, OTU-tables were rarefied to 1,000 reads. Alpha-diversity indices (Shannon index, Chao index and number of observed species) were calculated using Qiime (alpha_diversity.py). The significance of differences between samples from different compartments was tested using the Kruskal-Wallis test (krus.test in R, p<0.01). To estimate beta-diversity, OTU-tables were normalized using the cumulative-sum scaling (CSS) method (Paulson et al., 2013). Bray-Curtis distances between samples were used for principal coordinate analysis (PCOA, cmdscale function in R). To test the effect of location and compartment on the estimated explained variance, a PERMANOVA analysis was performed (Adonis function from vegan package, in R). Using the Bray-Curtis distance matrix as an input, the analysis was either constrained by site or by compartment. For each OTU the possible enrichment in one site and/or compartment was tested using a linear model (see Zgadzaj et al., 2016). The RA of all enriched OTUs was then summarized at the class level, separated by site and compartment.

### Microbial correlation network

To evaluate effects of compartment and site specificity, three networks were individually constructed for each kingdom. OTU tables for each dataset (bacterial, fungal, oomycetal) were restricted to OTUs that were present in at least two samples and comprised > 200 reads. For each table, Spearman correlation scores were calculated using the CoNet app (Faust et al., 2016) for Cytoscape (Shannon et al., 2003). Negative edges were discarded, and only edges with correlation scores of >0.6 were kept (p <0.05, Bonferroni corrected). For each node, affiliations to specific compartments and locations were estimated. If the sum of the normalized read count (using CSS) for one compartment or one site, respectively, comprised >50% of the read count, the affiliation was set to this compartment or location. Otherwise, the affiliation was set to the two dominant locations and compartments. Visualization was done with Cytoscape using the un-weighted force-directed layout. To estimate the mixing of nodes belonging to different affiliations we calculated a mixing parameter (Lancichinetti et al., 2009). To this end, the proportion of intergroup and intra-group edges (using the edge weights) was calculated for each of the occurring node affiliations To construct compartment-specific multi-kingdom co-occurrence networks, the OTU tables for the 16S and the two ITS datasets were restricted to samples from the respective compartments. Raw read count tables were merged to give one table per compartment. OTUs that appeared in less than ten samples were removed. These filtered multi-kingdom tables were used as an input for SparCC (Friedman et al., 2012). The analysis was conducted with default parameters and 100 bootstrap samples were used to infer pseudo-p.values. The inferred correlations were restricted to those having correlations >0.6 and <-0.6 (p<0.05, two-sided). Within the networks, proportions of inter- and intra-kingdom edges were calculated. Intra-kingdom refers to edges within bacterial OTUs, fungal OTUs or oomycetal OTUs, whereas inter-kingdom refer to edges between these groups. To estimate the strength of the antagonistic correlation between bacterial and fungal OTUs, a subset of those OTUs involved in negative correlation between the two groups was chosen. In this subset, for each OTU the number of negative bacterial-fungal correlations and the betweenness-centrality was calculated. The cumulative bacterial-fungal correlation refers to the sum of all inter-kingdom correlations for each fungal and bacterial OTU. OTUs belonging to taxonomic groups with less than five members were not shown. Visualization of the networks was done with Cytoscape (spring-embedded layout spring strength=5, spring rest length=15). To estimate the robustness of our SparCC-inferred network, we compared it to a network inferred from Spearman rank correlations (p>0.05, correlation strength >0.5 and <-0.5). OTU tables for the three kingdoms were filtered as described above and raw read counts were transformed to relative abundances (separately for each kingdom).

### Establishment of root-derived fungal and oomycetal culture collections

We grew *A. thaliana* Col-0 and *A. thaliana* relatives in Cologne Agricultural Soil (CAS) under greenhouse conditions. Thirty-six *A. thaliana* or *A. thaliana* relative *(Arabis alpina, Cardamine hirsuta)* individuals were harvested at three developmental stages (rosette, bolting and flowering stages). Similarly, two to six *A. thaliana* individuals from Geyen, Pulheim and Saint-Dié sites were dug out with their surrounding soil, transferred to sterile falcons and transported on ice to the laboratory. Plant roots were washed in sterile water to eliminate soil particles from the root surface and further three times in sterile water with shaking. In order to enrich for endophytic filamentous eukaryotes, roots were surface-sterilized for 1 min in 80% EtOH, followed by a second sterilization step for 1 min in 0.25% NaClO. The efficiency of the surface sterilization step was validated by printing the roots on TSB media. The emergence of hyphae from surface-sterilized root fragments was checked daily over two weeks and the corresponding microbial isolates where transferred to plates supplemented with antibiotics (Rimf^100^ Strp^100^ Amp^50^ Kn^50^ Tc^20^). DNA Isolation was performed using the DNAeasy Plant Mini Kit (Qiagen, Hilden, Germany) and fungal and oomycetal ITS regions were amplified by PCR using the ITS1F-ITS4 (fungi) and ITSO-5.8sORev (oomycetes) primer (94 °C for 2 min; 35 cycles of 94 °C for 30 sec, 53 °C for 30 sec and 72 °C for 1.5 min, 10 min at 72 °C).

### Culture collection comparison to natural sites

To compare the diversity of the fungal and oomycetal isolates retrieved by culturing, Sanger sequences were compared to amplicon sequences from natural sites (Geyen, Pulheim, Staint-Dié and CAS). ITSx (Bengtsson-palme et al., 2013) was used to extract the ITS region from all Sanger sequences, which were mapped against the representative OTU sequences (from all OTUs appearing in Root samples) from the respective datasets (Fungal, Oomycetal, using usearch_global) at a 97% sequence identity threshold. Recovery rates were defined as the number of recovered root-associated OTUs at a cumulative RA of 60% and 80%, respectively. To compare the taxonomic diversity between the culture collection and the root-associated OTUs, the amount of representative isolates was related to the number of core root-associated OTUs. These OTUs were defined as OTUs that appear within at least one location (including CAS samples) with more than 0.1% RA within all root samples from this site. OTUs with weak taxonomic assignment at the class level were excluded (RDP bootstrap >0.5). Sanger-derived sequences for all isolates were compared at a 100% identity threshold (using usearch – usearch -global) and isolates that shared 100% sequence identity and that were isolated at the same site were used for the construction of phylogenetic trees. The representative sequences were aligned using the MAFFT webservice (G-INS-i option and an un-align level of 0.4; Katoh et al., 2017). Maximum-likelihood trees were inferred with the RAxML webservice (Boc et al., 2012), using the default parameters. Trees were visualized using iTol (Letunic et al., 2016).

### Microbiota reconstitution experiments in the gnotobiotic FlowPot system

Bacterial strains were cultivated from their glycerol stock (pellets of bacterial colonies stored in 50% glycerol at -80 °C) in 96-deep-well plates containing 400 μl of 50% TSB (Tryptic Soy Broth, Sigma) for six days at 25 °C and subsequently pooled (in equal ratios). This bacterial pool was centrifuged at 4,000 xg for 10 min and re-suspended in 10 mM MgCl_2_ to remove residual media and bacteria-derived metabolites. Prior to inoculation, OD_600_ was adjusted to 0.02 (10^7^ cells/ml). Fungal strains were cultivated from their glycerol stocks (pieces of fungal mycelium stored in 30% glycerol stock at -80 °C) individually in PGA (Potato Glucose Agar, Sigma-Aldrich) including antibiotics (see above) for seven days and re-transferred to PGA for another seven days. Pieces of mycelium were harvested using sterile tips into 1 ml of MgCl_2_ containing one stainless steel bead (3.2 mm) and subsequently crushed with a paint shaker (SK450, Fast & Fluid Management, Sassenheim, Netherlands) for 10 min and pooled in equal ratios. Oomycetal strains were cultured in PGA for seven days and treated as described for fungal strains. Preparation of the gnotobiotic FlowPot system was carried out as previously described^16^. Microbial mixture was adjusted at a biomass ratio of 4:1 (eukaryotes:prokaryotes, as assessed by Joergensen et al., 2006), using 1mL (10^7^ cells) bacteria inoculum and 50μL of both fungal and oomycetes inocula (2.5 mg each) in 50mL ½ MS (Murashige + Skoog Medium including Vitamins, Duchefa), which were then inoculated in the FlowPot using a 50mL syringe. Prior inoculation, *A. thaliana* Col-0 seeds were sterilized and stratified for 4 days in the dark at 4°C. Between 4 to 5 hours after microbial inoculation, 10 seeds were sown per pot and the closed boxes incubated at 21°C, 10 hours light (intensity 4) and at 19°C, 14 hours dark for four weeks. After incubation, plant shoots were cut and weighted for fresh weight assessment. To account for experiment-to-experiment variation, raw shoot fresh weight values were normalized to the average shoot fresh weight of control plants (Microbe Free) per biological replicate and then all replicates were compared together, using *ggplot2* and *agricolae* packages in R. Kruskal-Wallis and Dunn’s *post-hoc* tests were used to test for significant differences. In the bacterial depletion experiments, bacterial rescue of plant growth was calculated by comparing the growth in shoot fresh weight of fungi-and-bacteria-inoculated plants relative to the bacteria-only inoculated plants (data not shown) (e.g: (BF-Comm/MF)/(B-Comm/MF)=Relative bacterial plant growth rescue). For community profiling, matrix samples were harvested and roots were thoroughly washed in water, dried on sterilized Whatman glass microfiber filters (GE Healthcare Life Sciences), transferred into lysing matrix E tubes (MP Biomedicals), frozen in liquid nitrogen and stored at -80 °C. DNA extraction and amplicon sequencing was performed as described above. The microbiota reconstitution experiment includes three fully independent biological replicates, each containing three technical replicates.

### Direct mapping for synthetic communities and downstream analysis

The reads of the synthetic communities were processed as described before for the respective barcodes until demultiplexed and filtered raw reads were available. Raw reads where directly mapped to the reference sequences for the respective communities (using the usearch-global command from usearch, sequence identity threshold of 97%). OTU-tables were inferred from these mapped reads. All alpha and beta diversity indices were calculated as described before. Relative abundance plots were produced from relative OTU counts (RA%) per sample and plotted using R. Raw OTU tables were rarefied with a subsampling depth of 1,000 reads using qiime (single_rarefaction.py) and the alpha-diversity indices were calculated using alpha_diversity.py. Alpha-diversity values (observed OTUs) were then plotted using ggplot in R and corrected for experiment-to-experiment variation. Beta diversity was estimated as described before. Permutational multivariate analyses of variance were performed in R using the function capscale. Statistical significance of the ordinations as well as confidence intervals for the variance was determined by an ANOVA-like permutation test (functions permutest and anova.cca) with 999 permutations (**Table S5**). To identify strains enriched in single-microbial class inoculation conditions (B, F, O) compared to combined-microbial class inoculation conditions (BF, BO, ALL), we employed linear statistics on RA values (log2, > 5 %o threshold) using a script described previously (Bulgarelli et al., 2012). Ternary plots were constructed as previously described (Bulgarelli et al., 2012). To visualize the distance between clusters of the same microbial inoculation condition, average Bray-Curtis distances were calculated per biological replicate and normalized to control cluster (e.g: (B-BF/B-B)) (**Figure S10**). Kruskal-Wallis and Dunn’s *post-hoc* tests were used to determine significant differences. For the comparison between natural communities and SynComs (**Figure S17**), amplicon sequencing data derived from the sequencing from the three natural sites (Pulheim, Geyen and Saint-Dié) and from plants grown in Cologne Agricultural Soil (CAS) were processed together with the sequencing data derived from the reconstitution experiment. All reads for the respective datasets were pooled and processed as described in the 16s and ITS read processing sections. Assuming that our culture collections contained the most abundant OTUs from *A. thaliana* roots, we used only the 100 most abundant root associated OTUs for calculating Bray-Curtis distances between samples.

### High-throughput bacterial-fungal interaction screen

The protocol was adapted from Figueroa-López et al., 2014. The screen was performed in triplicate and independently validated by another random screening (**Figure S13**). Spores from three weeks-old fungal cultures were re-suspended in 10mL sterile water, transferred in a 12mL falcon tube, and centrifuged 3 minutes 2,000 rpm. After three washes with sterile water, spore concentration was adjusted to 1×10^6^ spores/mL in TSB20% and the solution was kept at 4°C. Bacterial cells were grown in 96-well plates in TSB 20% for 6 days to reach the stationary phase. After a centrifugation step (20 minutes 4,000 rpm at 4°C), the TSB medium was removed and replaced by a fresh medium (200 μl in each well). Two 96 well-optical bottom plates (ThermoFisher Scientific/Nunc, Rochester, USA) were used for the screening. Plate A corresponds to the screening plate and contained fungal spores in interaction with bacteria or fungal spores alone. Plate B corresponds to a control plate inoculated with bacteria only to assess bacterial OD, as well as bacterial autofluoresence intensity. Plate A was filled with 150 μl of TSB20% medium or 160 μl of TSB20% medium for the control without bacteria and plate B was filled with 190 μl of TSB20% medium. Then, 40 μl of fungal spores (1×10^6^ spores/mL) were disposed in all wells of the plate A and 10 μl of 6 days old bacterial culture in each well of both plates, except for the control wells containing the fungus alone in plate A. After 48 hours of incubation at 25°C, the bacterial cells from plates A and B were washed away with a multichannel pipet and the wells were further washed two times with 200 μl of PBS1x and incubated overnight at 4°C (dark) in 100 μl of PBS1x supplemented with 1μg/mL of WGA Alexa fluor 488 conjugated (stock solution: 1 mg/mL, Invitrogen/Molecular Probes, Eugene, USA). The solution was washed away and the wells were further washed two times in 200 μl of PBS1x and finally resuspended in 200 μl of PBS 1x. The fluorescence intensity, reflecting fungal growth in each well is measured using a plate reader (Excitation/Emission 490/530, Tecan Infinite 200 PRO, Männedorf, Switzerland). Log2 transformed relative fluorescence values, reflecting the ability of a specific bacterium to restrict or promote fungal growth in the presence versus absence of bacterial competitors were calculated as shown in **Figure S13**. The Bacterial phylogenetic tree was constructed based on the full sequences of bacterial 16S rRNA. Sequences were aligned with MUSCLE using default parameters and MEGA 5.0 was used to construct a Neighbor Joining tree. The bootstrap consensus tree inferred from 1,000 replicates was edited in iTOL (Letunic et al., 2016).

## Quantification and Statistical analysis

No statistical methods were used to pre-determined sample sizes. Data collection and analysis were performed blinded to conditions in all experiments. A p-value of less than 0.05 was considered as significant.

## Data and software Availability

All scripts for computational analysis and corresponding raw data are available at https://github.com/ththi/Microbial-Interkingdom-Suppl.

## Author contributions

S.H. and P.S.-L. initiated, coordinated and supervised the project. S.H. and P.D. collected root material and performed culture-independent community profiling. S.H. isolated root-associated fungi and oomycetes. TT analysed culture-independent 16S rRNA and ITS amplicon sequencing data. R.G.-O. contributed to bioinformatics tools. M.A. and E.K. identified *A. thaliana* populations. P.D. performed all microbiota reconstitution experiments in the FlowPot system. T.T. and P.D. analysed the recolonization data. S.H. developed, performed and analysed the high-throughput bacterial-fungal interaction screen. P.D., T.T., P.S.-L. and S.H. wrote the manuscript.

## Acknowledgements

We thank J Kremer and SY He for sharing the FlowPot utilization protocol before publication. We thank Neysan Donnelly for scientific English editing. This work was supported by funds to S.H from a European Research Council starting grant (MICRORULES) and funds to P.S.-L. from the Max Planck Society, a European Research Council advanced grant (ROOTMICROBIOTA) and the ‘Cluster of Excellence on Plant Sciences’ program funded by the Deutsche Forschungsgemeinschaft.

## Accession numbers

Sequencing reads from natural site samples and microbiota reconstitution experiments (MiSeq 16S rRNA and ITS reads) have been deposited in the European Nucleotide Archive (ENA) under accession numbers PRJEB27146 and PRJEB27147, respectively.

## Supplementary Figures

**Figure S1:**
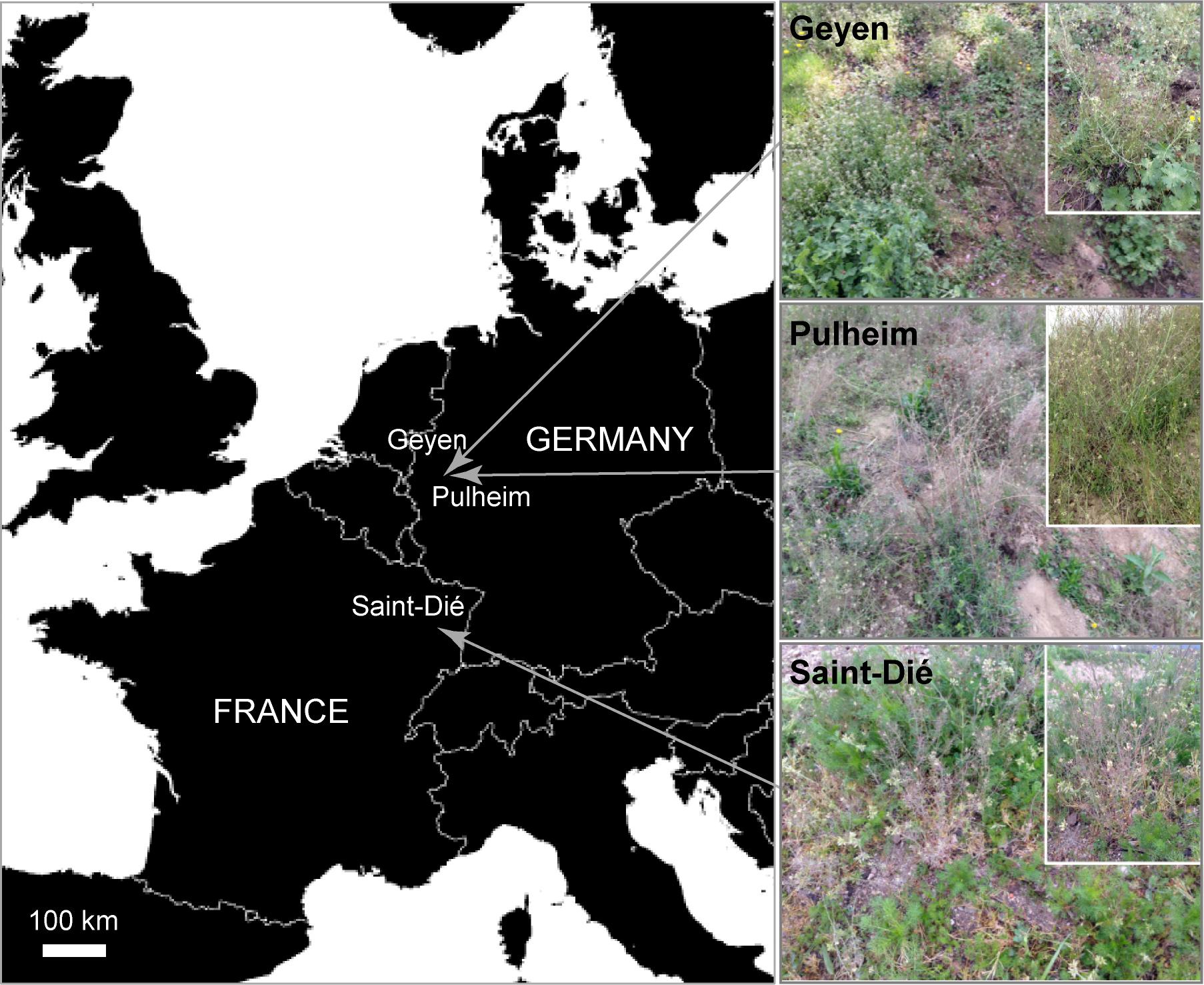
Location of the three natural *A. thaliana* populations and plant developmental stages. The sites include two geographically-related populations in Geyen and Pulheim (Germany), and a more distant population in Saint-Dié (France). All plants were harvested in spring 2014 at the flowering stage.

**Figure S2:**
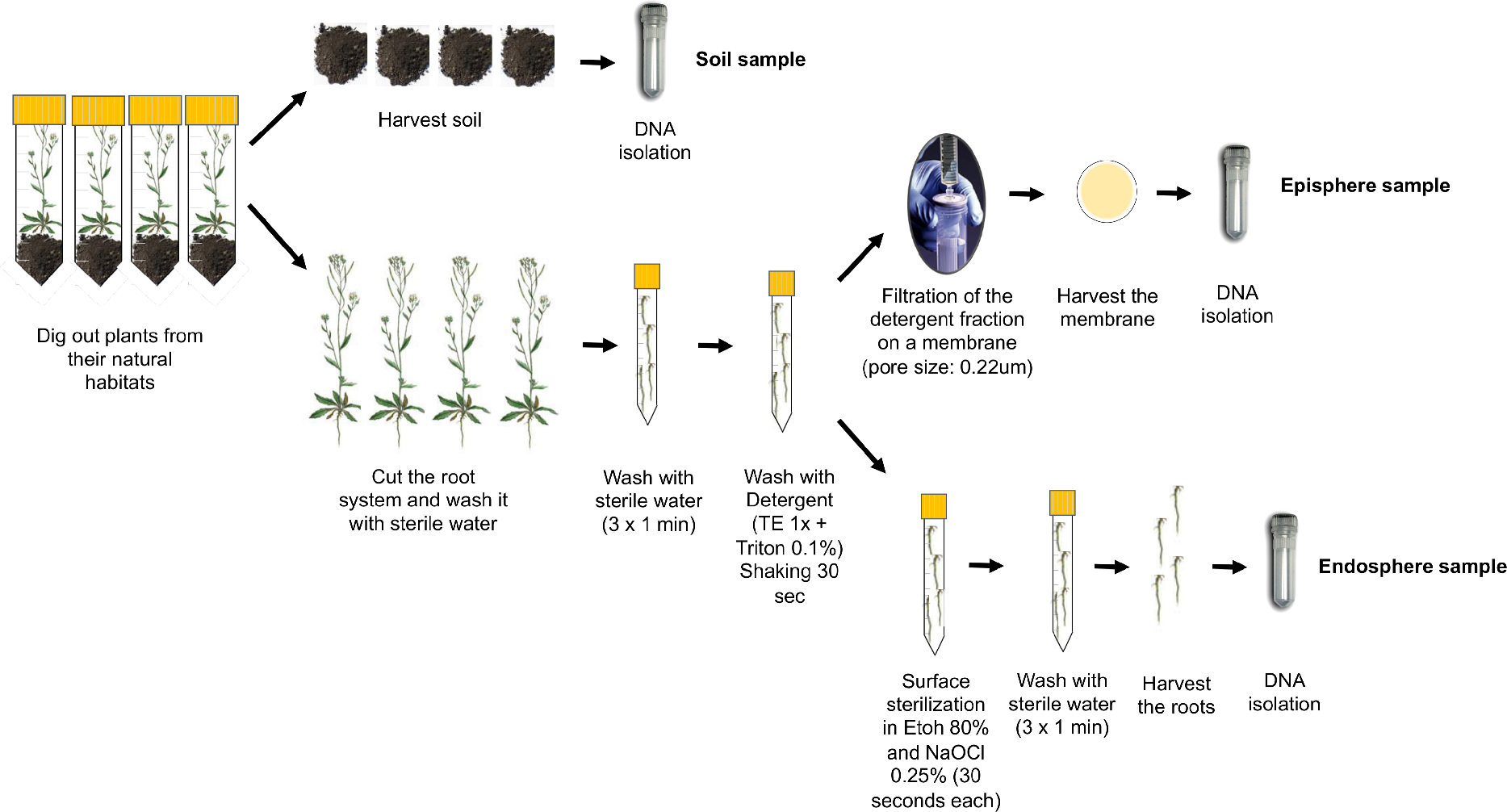
Fractionation protocol. For each biological replicate (n=4), four plant individuals were dug out from their natural habitats and transferred to 50-mL falcons on ice in the laboratory. Soil particles not in direct contact with the root system were harvested and pooled (soil sample). The plants were harvested and the four root systems were cut and washed briefly in sterile water. After three washes in sterile water, microbes were detached from the root surface using a detergent-based method (shaking for 30 seconds). The detergent fraction (containing episphere microbes) was then filtered on a 0.22-μM pore size membrane (episphere sample). The cleaned root systems were surface-sterilized, treated with bleach and washed three times in sterile water (endosphere sample).

**Figure S3:**
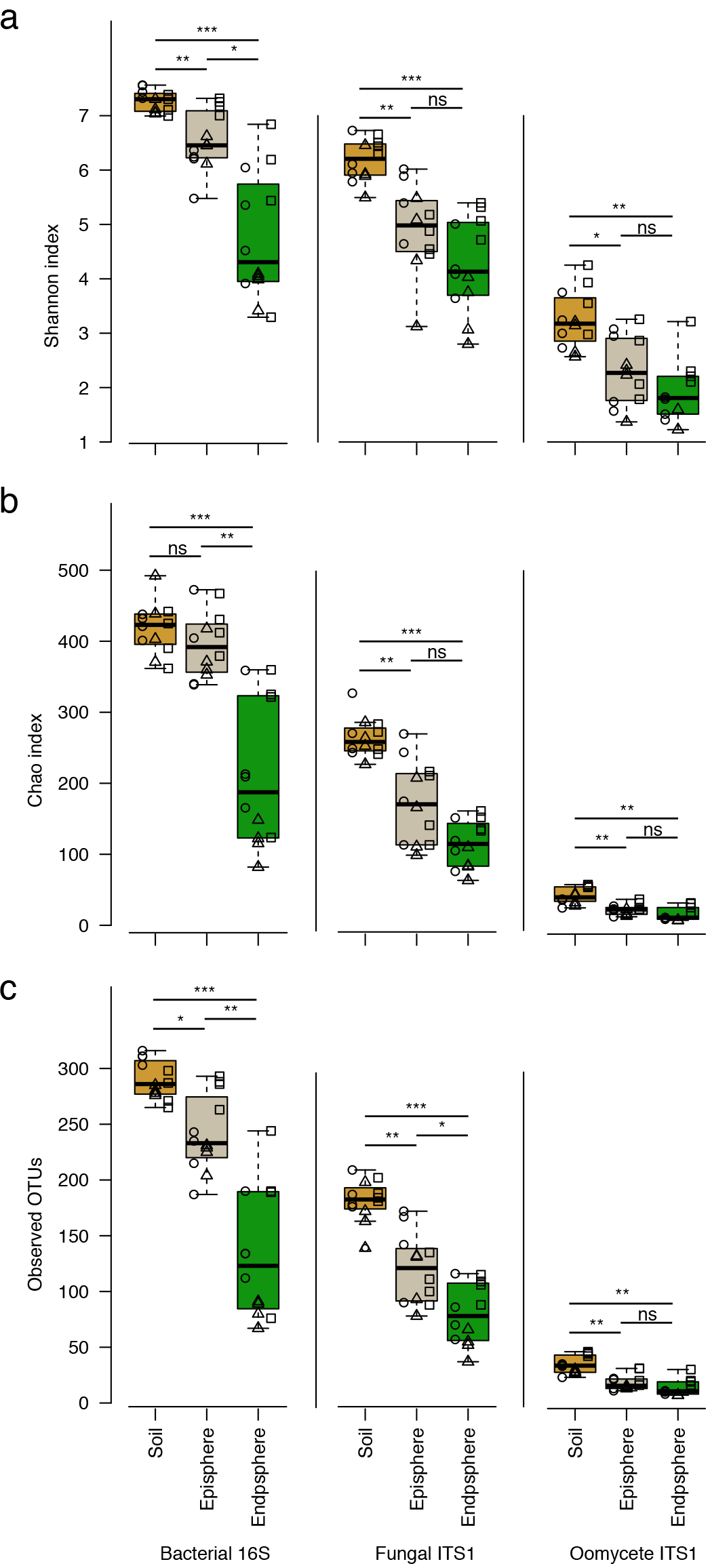
Microbial alpha diversity across sites and compartments. **a,** Boxplot for the Shannon index, **b,** Boxplot for the Chao index and **c,** Boxplot for the Observed OTUs (left: bacteria, middle: fungi, right: oomycetes). For each of the three indices, all samples from a given site are taken into account (rarefied to 1,000 reads). Individual data points within each box correspond to samples from the three natural sites (circles = Geyen, triangles = Pulheim, squares = Saint-Dié). ns = not significant, * = *p*<0.01, ** =*p*<0.001,*** =*p*<0.0001 (Kruskal-Wallis test)

**Figure S4:**
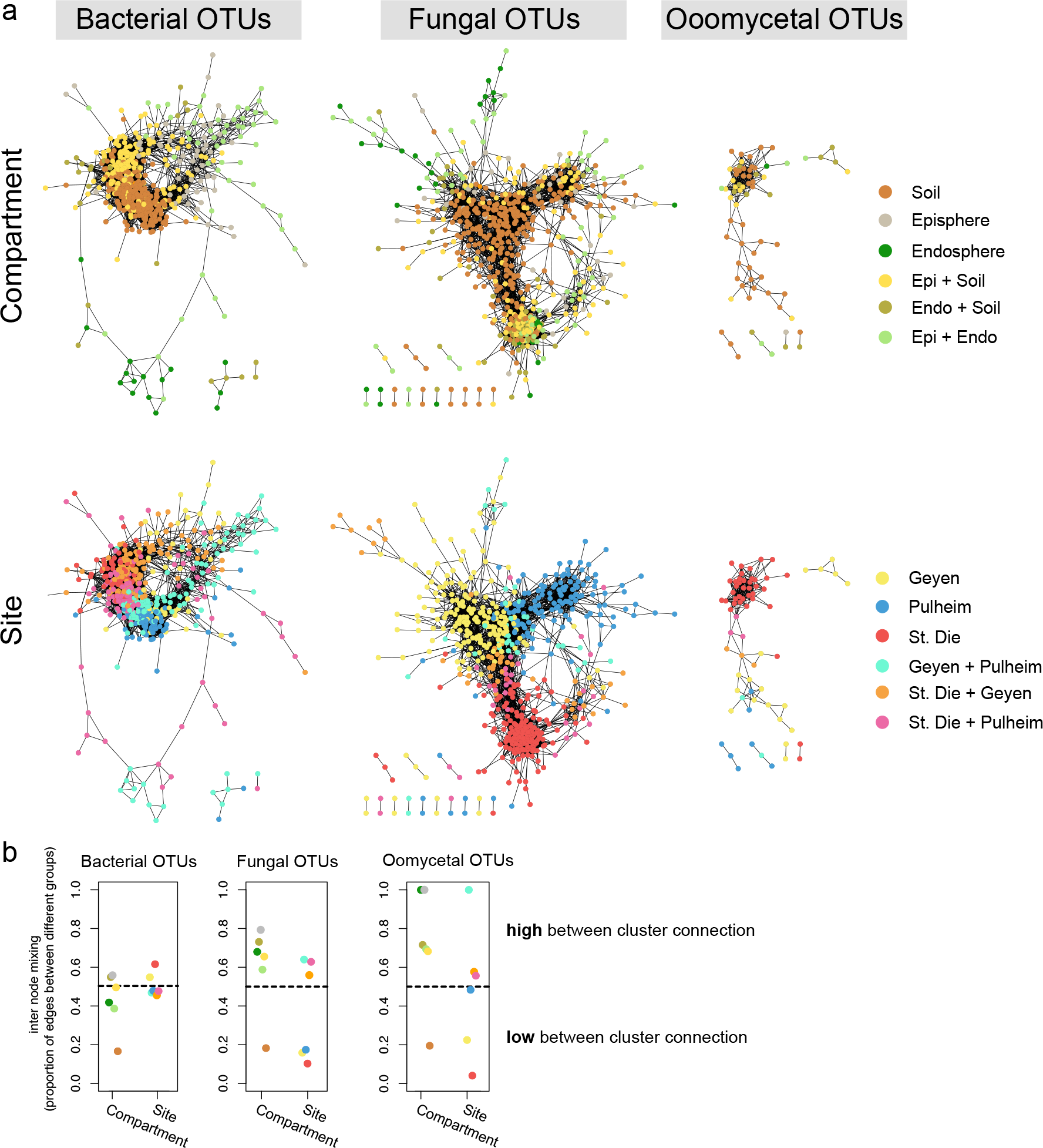
Microbial co-occurrence networks across sites and compartments. **a,** Networks showing microbial OTU co-occurrence patterns across compartments and sites for individual microbial groups. Spearman correlation-based networks are shown for bacterial (left), fungal (middle) and oomycetal (right) OTUs. Note that microbial OTUs with >200 reads and that were present in at least two samples were considered and that only edges with a correlation score >0.6 were kept (p <0.05, Bonferroni corrected). For each microbial OTU, the compartmental (upper part) and locational (lower part) affiliation is indicated (>50% reads coming from one or a combination of two compartments or sites, respectively). **b,** Internode mixing for each microbial network. For each microbial network (left: bacteria, middle: fungi, right: oomycetes), the amount of internode mixing is plotted considering the compartmental or locational affiliation of each OTU.

**Figure S5:**
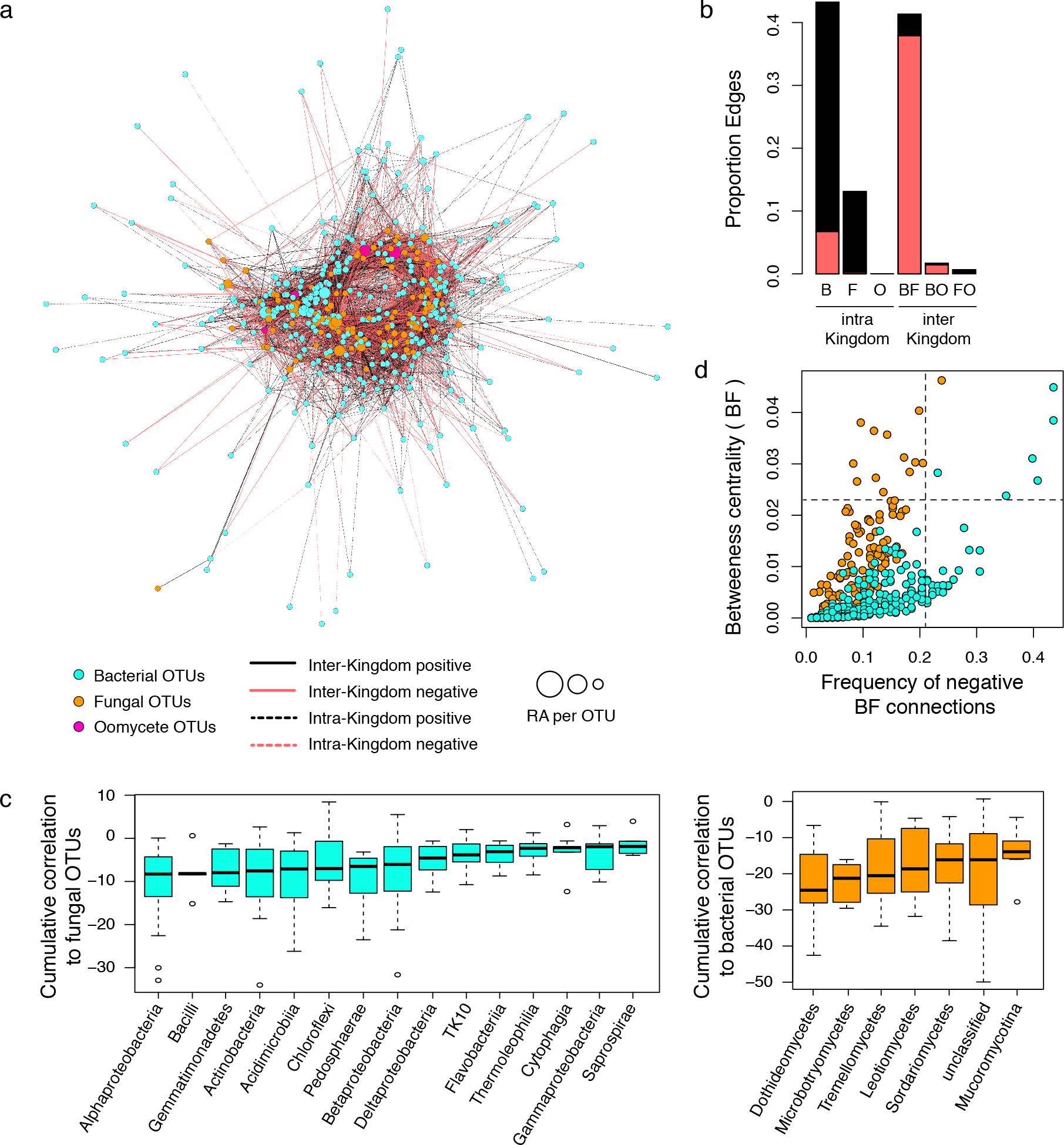
Microbial network of the *A. thaliana* episphere microbiota. **a,** Correlation-based network of microbial epishere-associated OTUs detected in three natural *A thaliana* populations (Pulheim, Geyen, Saint-Dié). Each node corresponds to a microbial OTU and edges between nodes correspond to either positive (black) or negative (red) correlations inferred from OTU abundance profiles using the SparCC method (pseudo *p*-value >0.05, correlation values <-0.6 or >0.6). OTUs belonging to the different microbial kingdoms have distinct colour codes and node size reflects their relative abundance in the episphere compartment. Intra-kingdom correlations are represented with dotted lines and inter-kingdom correlations by solid lines. **b,** Proportion of edges showing positive (black) or negative (red) correlations in the microbial episphere network. B: bacteria, F: fungi, O: oomycetes. **c,** Cumulative correlation scores measured in the microbial network between bacterial and fungal OTUs. Bacterial (left) and fungal (right) OTUs were grouped at the class level (> five OTUs/class) and sorted according to their cumulative correlation scores with fungal and bacterial OTUs, respectively. **d,** Hub properties of negatively correlated bacterial and fungal OTUs. For each fungal and bacterial OTU, the frequency of negative inter-kingdom connections is plotted against the betweenness centrality inferred from all negative BF connections (cases in which a node lies on the shortest path between all pairs of other nodes). The five microbial OTUs that show a high frequency of negative inter-kingdom connections and betweenness centrality scores represent hubs of the “antagonistic” network and are highlighted with numbers.

**Figure S6:**
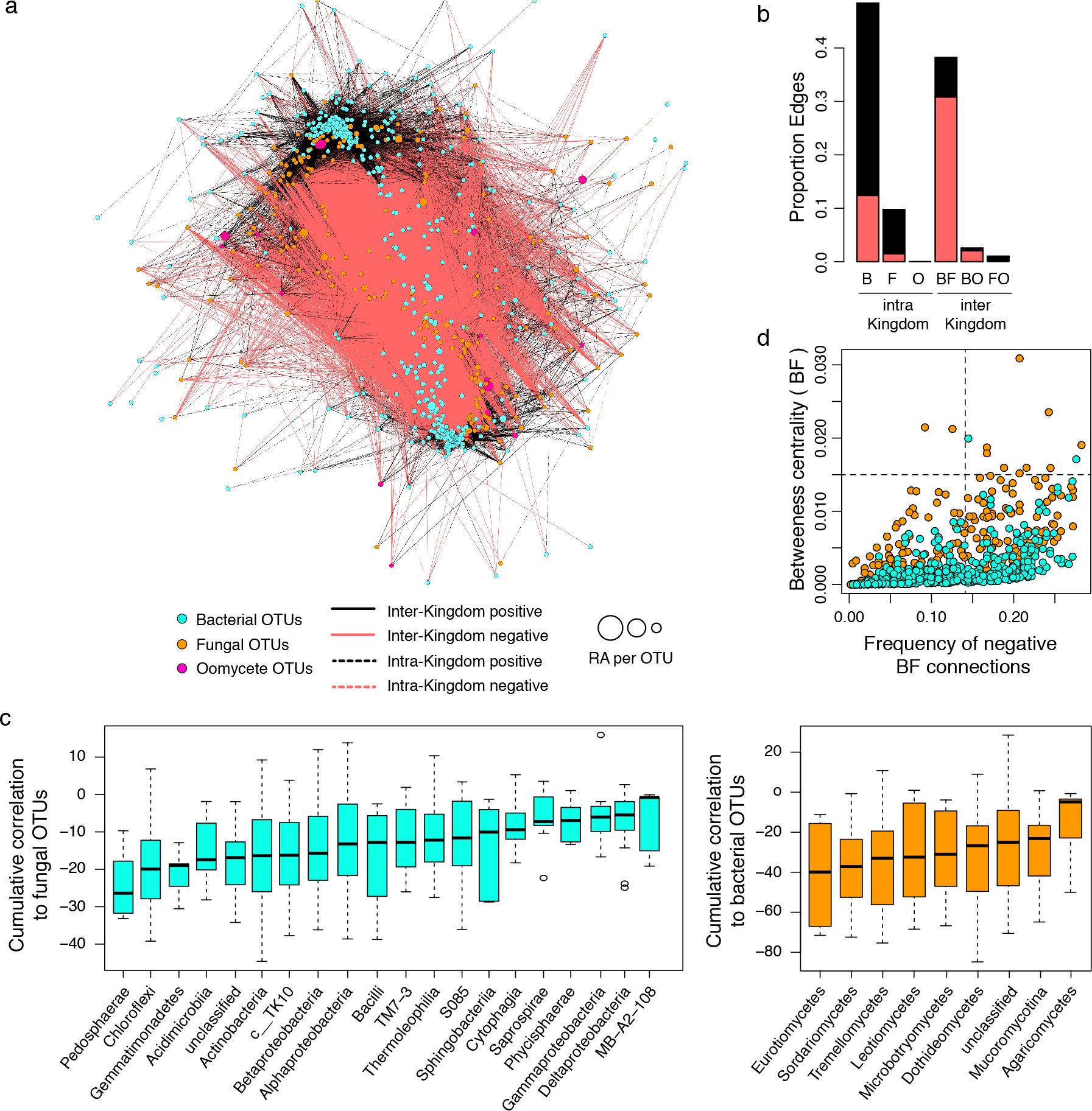
Microbial network of the soil microbiota. **a,** Correlation-based network of microbial soil OTUs detected in three natural sites with *A. thaliana* populations (Pulheim, Geyen, Saint-Dié). Each node corresponds to an OTU and edges between nodes correspond to either positive (black) or negative (red) correlations inferred from OTU abundance profiles using the SparCC method (pseudo *p*-value >0.05, correlation values <-0.6 or >0.6). OTUs belonging to different microbial kingdoms have distinct colour codes and node size reflects their relative abundance in the soil compartment. Intra-kingdom correlations are represented with dotted lines and inter-kingdom correlations by solid lines. **b,** Proportion of edges showing positive (black) or negative (red) correlations in the microbial soil network. B: bacteria, F: fungi, O: oomycetes. **c,** Cumulative correlation scores measured in the microbial network between bacterial and fungal OTUs. Bacterial (left) and fungal (right) OTUs were grouped at the class level (> five OTUs/class) and sorted according to their cumulative correlation scores with fungal and bacterial OTUs, respectively. **d,** Hub properties of negatively correlated bacterial and fungal OTUs. For each fungal and bacterial OTU, the frequency of negative inter-kingdom connections is plotted against the betweenness centrality inferred from all negative BF connections (cases in which a node lies on the shortest path between all pairs of other nodes). The five microbial OTUs that show a high frequency of negative inter-kingdom connections and betweenness centrality scores represent hubs of the “antagonistic” network and are highlighted with numbers.

**Figure S7.**
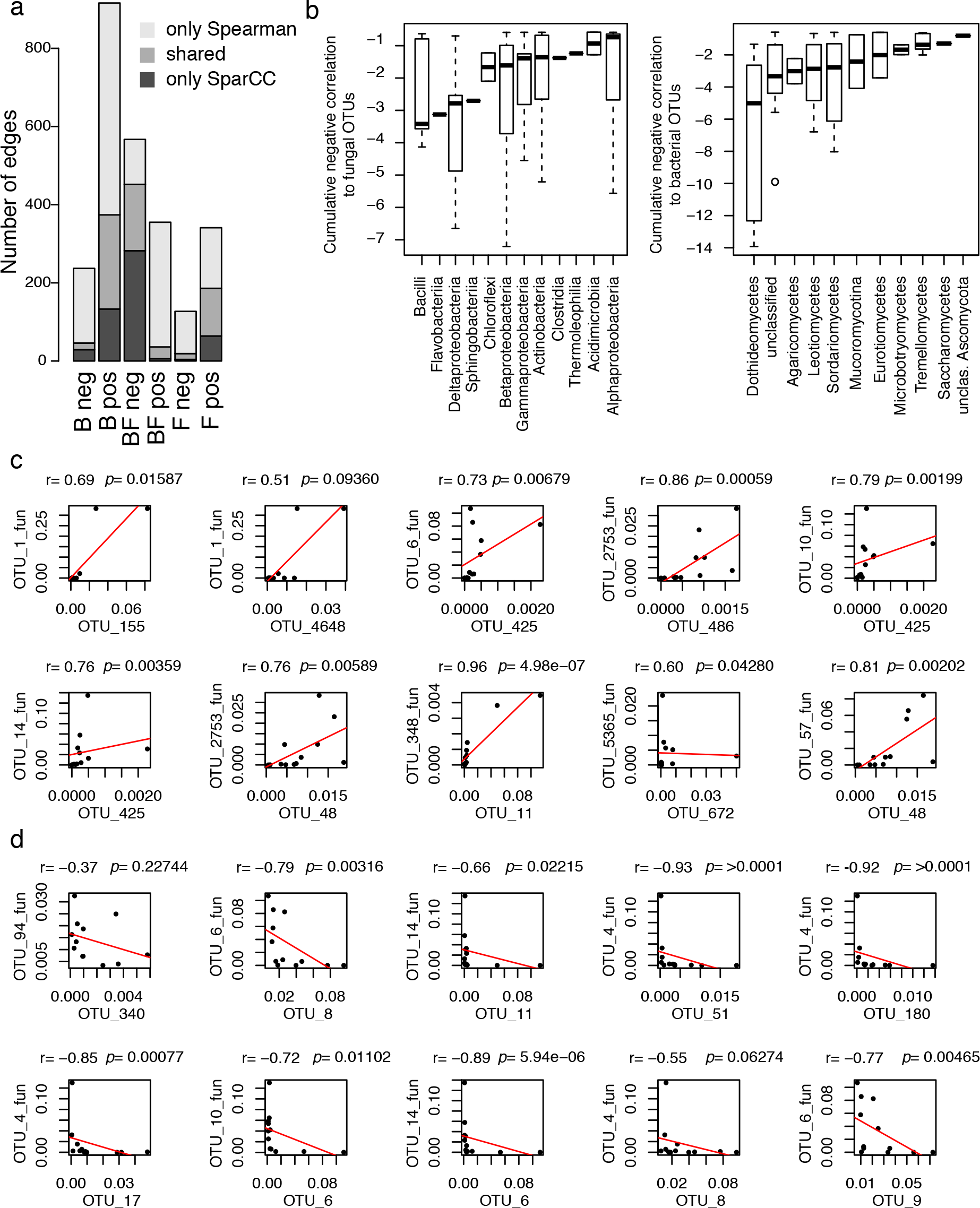
Validation of sparCC-defined (anti-)correlated OTUs using Spearman correlation. **a,** Number of edges that are either unique to one of the two networks or shared by the two (B = connections between bacterial OTUs, F = connections between fungal OTUs, BF = connections between fungal and bacterial OTUs, pos = positively correlated connections, neg = negatively correlated connections). **b,** Cumulative negative correlations across taxonomic groups inferred by Spearman correlation. **c,** The ten most positively correlated OTUs from the sparCC network and the corresponding correlation inferred by Spearman correlation based on the relative abundance of the OTUs. **d,** The ten most negatively correlated OTUs from the sparCC network and the corresponding correlation inferred by Spearman correlation based on the relative abundance of the OTUs.

**Figure S8:**
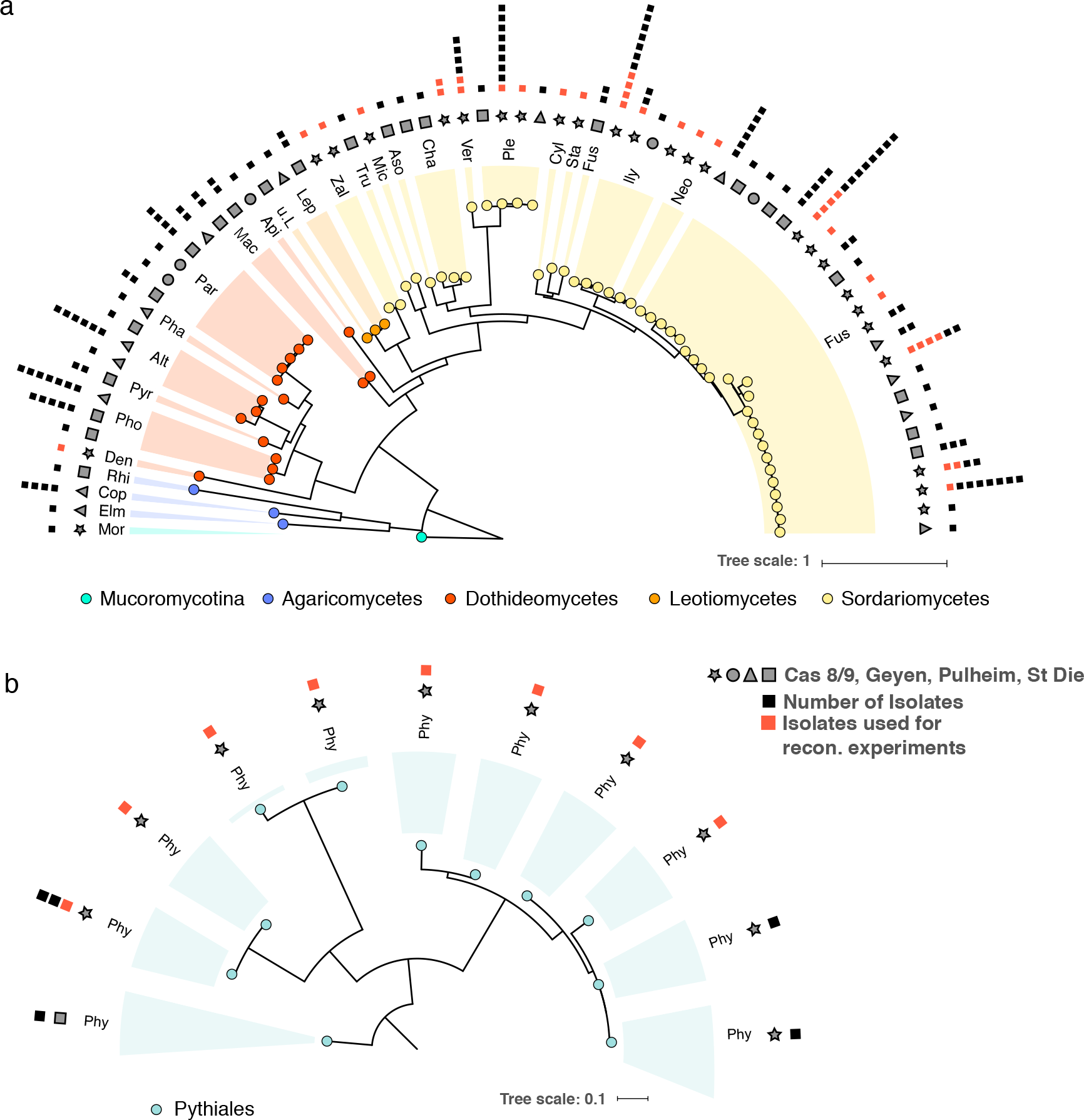
Phylogenetic diversity of fungal and oomycetal culture collections. Maximum likelihood trees of Sanger-sequenced ITS sequences for all isolates that show non identical ITS or that originate from different sites. **a,** Fungal isolates (Mor = *Mortierella*, Elm = *Elmerina*, Cop = *Coprinopsis*, Rhi = *Rhizoctonia*, Den = *Dendryphion*, Pho = *Phoma*, Pyr = *Pyrenochaeta*, Alt= *Alternaria*, Pha = *Phaseolina*, Par = *Paraphoma*, Mac = *Macrophomina*, Api = *Apiosporina*, u.L. = unclassified *Leotiomycete*, Lep = *Leptodontidium*, Zal = *Zalerion*, Tru = *Truncatella*, Mic = *Microdochium*, Aso = *Asordaria*, Cha = *Chaetomium*, Ver = *Verticillium*, Ple = *Plectosphaerella*, Cyl = *Cylindrocarpon*, Sta = *Stachybotrys*, Fus = *Fusarium*, Ily = *Ilyonectria*, Neo = *Neonectria).* **b,** Oomycetal isolates (Phy = Phytium). The first outer ring indicates the origin of each isolate, the second ring shows the number of isolates with 100% sequence identity that were isolated from the same site, therefore representing clonal duplicates. Isolates that were used in the reconstitution experiments are highlighted with red squares.

**Figure S9:**
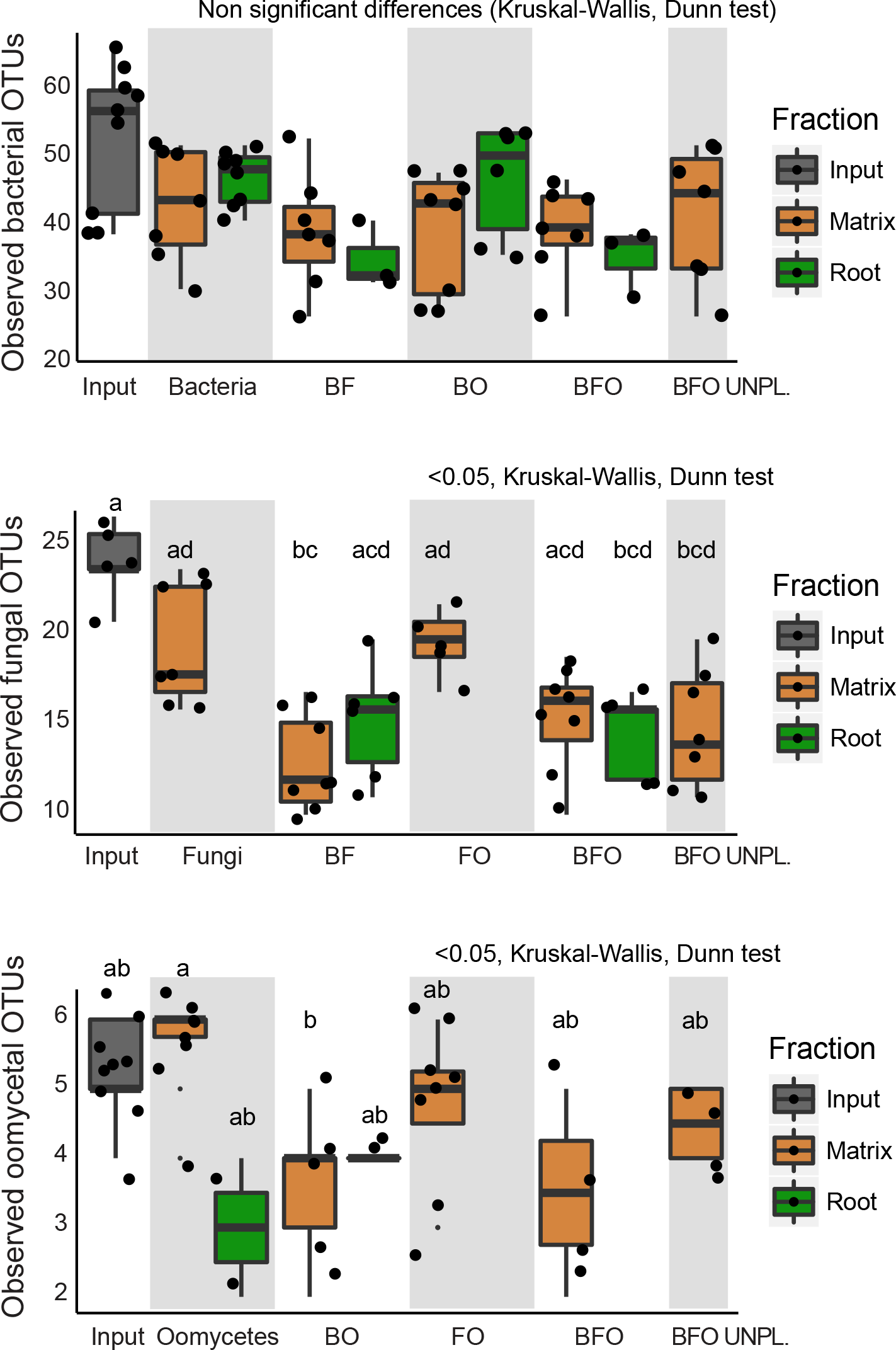
Microbial alpha diversity in matrix and root compartments in a multikingdom microbiota reconstitution system. Germ-free plants were re-colonized with root-derived bacterial (148), fungal (34) and oomycetal (9) isolates in the FlowPot system and matrix and root compartments were harvested after four weeks. Observed bacterial (upper panel), fungal (middle panel) and oomycetal (lower panel) OTUs in matrix (brown) and root (green) samples, as well as in the corresponding microbial input communities inoculated in the FlowPot system at T0 (grey) (p<0.05, Kruskal-Wallisa and Dunn’s *post-hoc* tests). Data points are missing for several root samples due to the absence of living plants in the corresponding treatment. B: bacteria, F: fungi, O: oomycetes, BO: bacteria and oomycetes, BF: bacteria and fungi, FO: fungi and oomycetes, BFO: bacteria, fungi and oomycetes, UNPL: unplanted pots. Note the significant decrease in observed fungal and oomycetal OTUs in the presence *versus* absence of bacteria.

**Figure S10:**
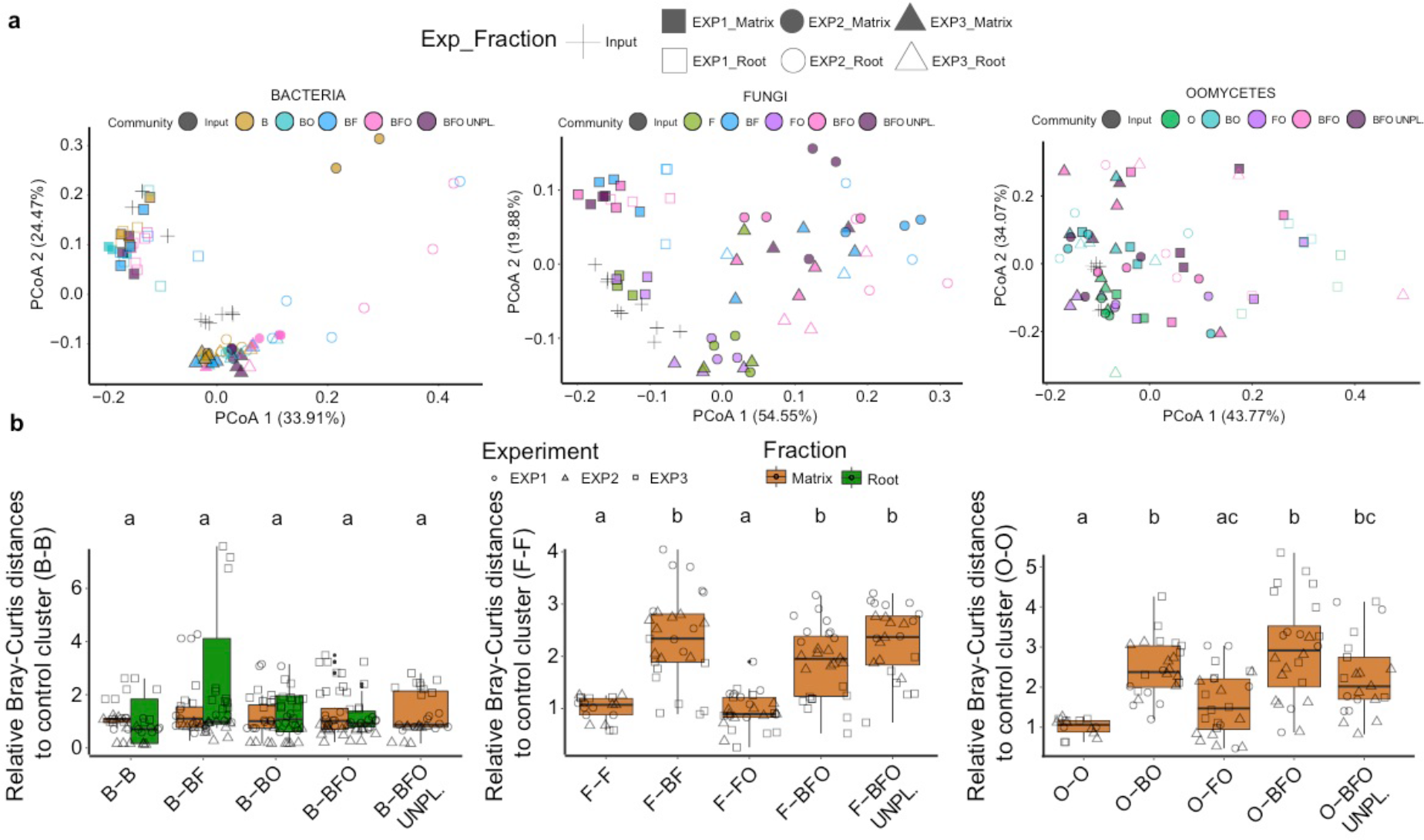
Microbial community structure in matrix and root compartments in a multi-kingdom microbiota reconstitution system. Germ-free plants were re-colonized with root-derived bacterial (148), fungal (34) and oomycetal (nine) isolates in the FlowPot system and matrix and root compartments were harvested after four weeks. **a,** PCoA plots of bacterial, fungal and oomycetal profiles (from left to right); shapes represent three biological replicates and colours depict different microbial combinations. B: bacteria, F: fungi, O: oomycetes, BO: bacteria and oomycetes, BF: bacteria and fungi, FO: fungi and oomycetes, BFO: bacteria, fungi and oomycetes, UNPL: unplanted pots. **b,** Relative Bray-Curtis distances between sample clusters of bacterial, fungal and oomycete profiles (from left to right) of matrix (brown) and root (green samples) to the control clusters (B-B, F-F and O-O) (i.e. the closer to 1, the more similar to the control cluster; see methods). Significant differences are depicted with different letters (Kruskal-Wallis and Dunn’s *post-hoc* tests, <0.05).

**Figure S11:**
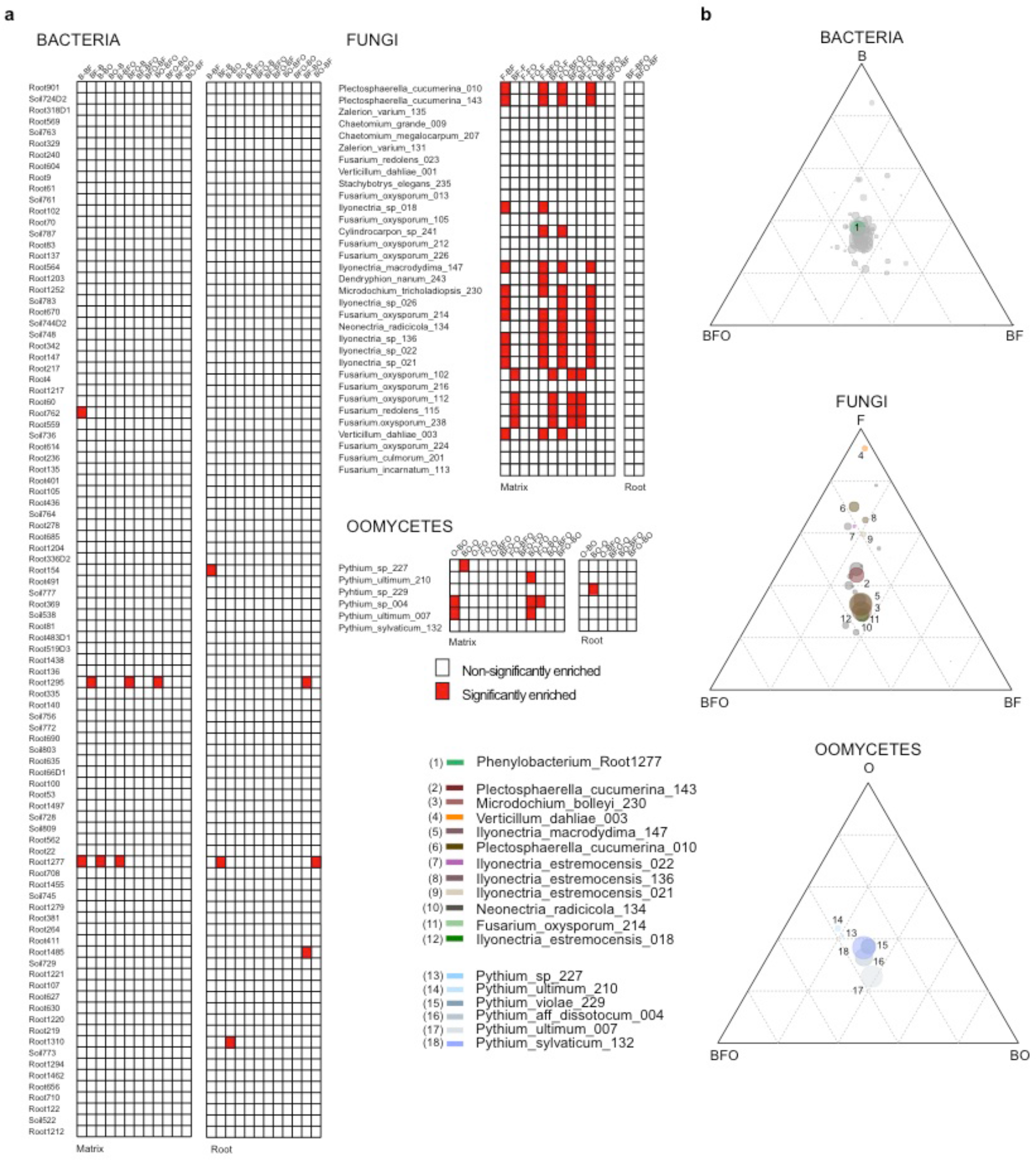
Effect of co-inoculation on microbial strain enrichment. **a,** Pairwise enrichment tests for bacterial, fungal and oomycetal strains in gnotobiotic experiments (Generalized Linear Model, p.adj.method=FDR, p-value<0.05) co-inoculated in different combinations (B: bacteria only, F: fungi only, O: oomycetes only, BO: bacteria and oomycetes, BF: bacteria and fungi, FO: fungi and oomycetes, BFO: full microbial community). Enriched strains in one combination compared to another one are depicted with a red block (e.g. Root762 is enriched in B compared to BF). **b,** Ternary plots representing the enriched strains (coloured circles) (Generalized linear model, p.adj.method=FDR <0.05) in each combination versus the other two combined. The size of the circles indicates the relative abundance of each strain and the closeness to each edge signifies a higher prevalence in that given condition.

**Figure S12:**
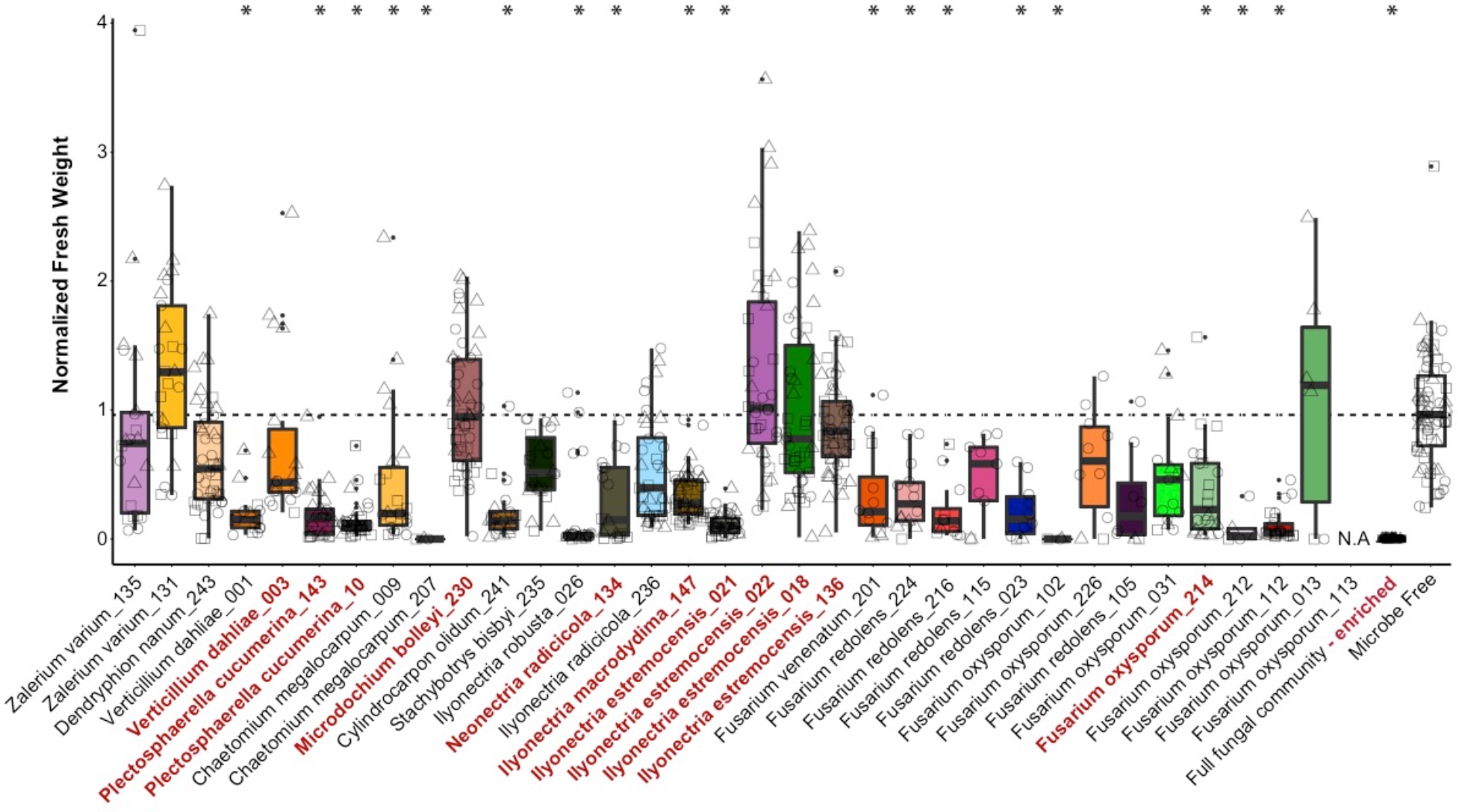
Effect of individual fungal isolates on *A. thaliana* growth in the FlowPot system. The boxplots depict normalized fresh weight of *A. thaliana* Col-0 plant shoots after three weeks of incubation with each of the 34 fungal strains used in the multi-kingdom microbiota reconstitution experiment (see **Figure 4**). For each boxplot, three biological replicates (depicted with different shapes) with at least three technical replicates are presented. Significant differences are indicated with an asterisk (p<0.01, Kruskal-Wallis, Dunn’s *post-hoc* tests). The 11 fungal strains enriched in the absence of bacteria (depicted in **Figure S11**) are highlighted in red and a 23-member fungal community lacking these 11 isolates remains deleterious for plant growth (fungal community-enriched). N.A.: data not available

**Figure S13:**
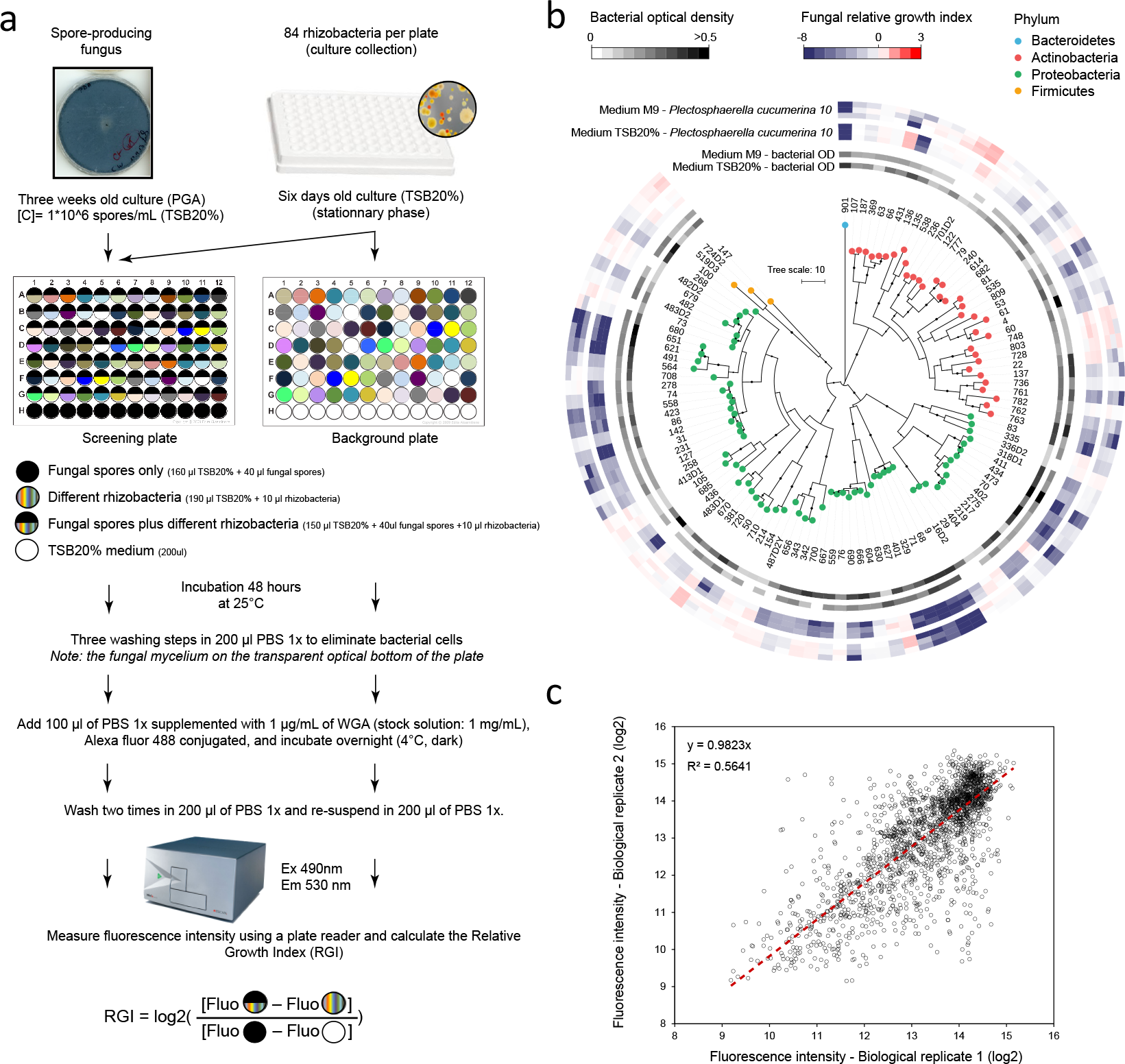
High-throughput fungal-bacterial interaction screen. **a,** Schematic overview of the experimental protocol. Fungal spores were equally distributed to the wells of a transparent-bottom 96 well plate and incubated in the presence or absence of different rhizobacteria (stationary phase) in liquid medium (screening plate). Bacteria were also grown in the absence of fungal spores as a control (background plate). After 48 hours of interaction, three washing steps were used to eliminate bacterial cells in suspension. Note that the fungal mycelium stick to the transparent optical bottom of the plate. After overnight incubation in Wheat Germ Agglutinin and two additional washes, the fluorescence intensity (reflecting fungal growth) was measured using a plate reader. The relative growth index was calculated as illustrated (see methods). **b,** Alteration of the growth of *Plectosphaerella cucumerina isolate 10* upon competition with phylogenetically diverse members of the bacterial root microbiota in minimum medium (M9) and a carbon-rich medium (20% TSB). The phylogenetic tree was constructed based on the full bacterial 16S rRNA gene sequences and bootstrap values are depicted with black circles. The heatmap depicts the log2 fungal relative growth index (presence vs. absence of bacterial competitors) measured by fluorescence (see above). Note the overall similar inhibitory activities in minimum and complex media. **c,** Validation experiment, in which the bacterial strains were re-screened against ten randomly selected fungi. Fluorescence intensities (log2) were compared with those obtained from the first biological replicate (y=0.9823, R^2^=0.5641).

**Figure S14:**
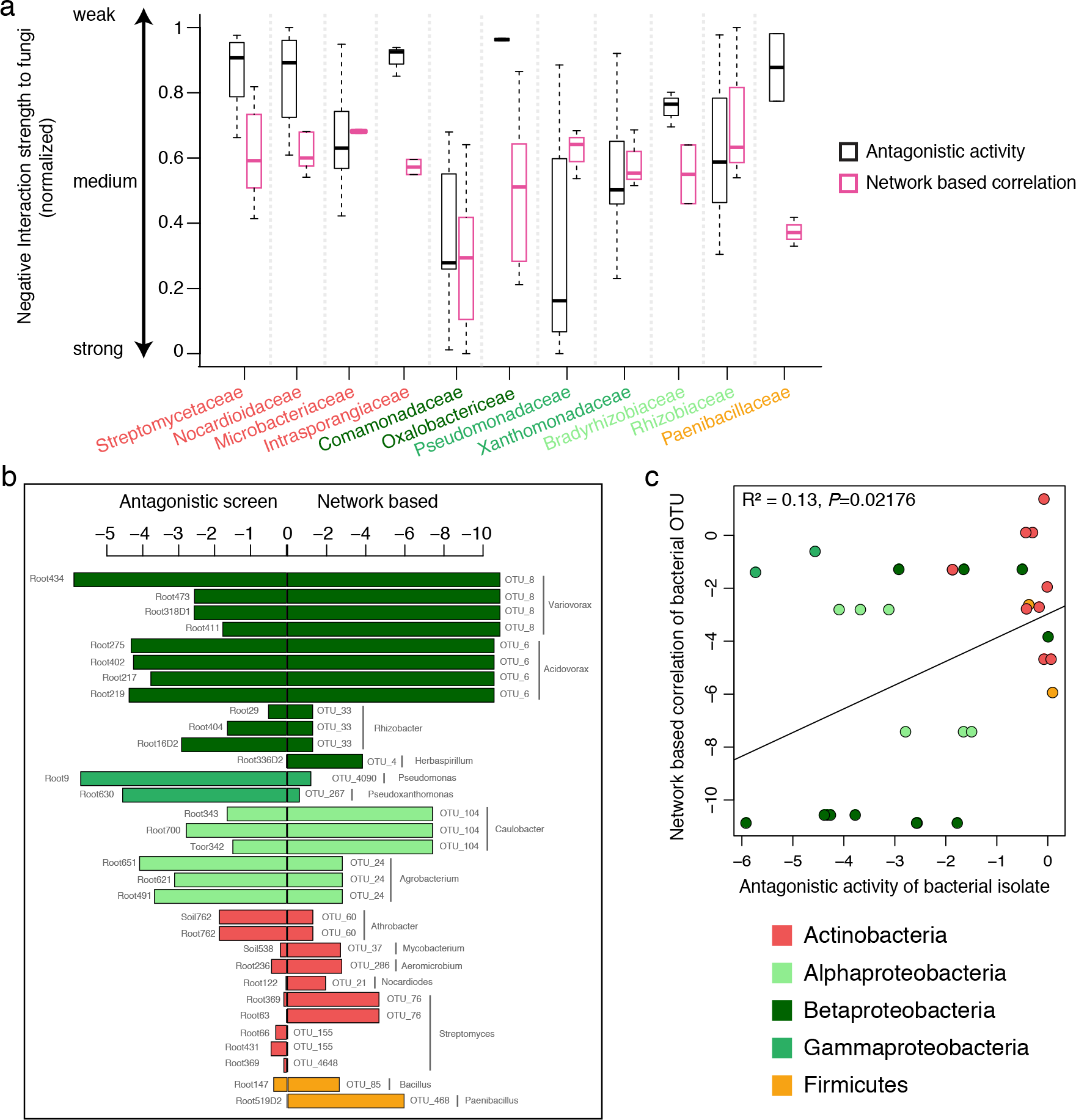
Comparison of network-derived correlations and experimentally tested interactions of bacterial families with fungal species. **a,** For each bacterial family shared between antagonistic screening and root-associated OTU network analysis, the average antagonistic activity against fungal isolates and the cumulative correlation to fungal OTUs in the network are represented. Bacterial families with more than two members were considered and values for both measurements were normalized to be in the same range. **b,** Direct comparison of bacterial OTUs from the root network with bacterial isolates used in the antagonistic screening. Each data point corresponds to a bacterial OTU-isolate pair (>97% sequence similarity, only best matching hits are shown). For each pair, the network-derived correlation with fungal OTUs (from the bacterial OTU) is plotted against the result from the antagonistic screening (from the bacterial isolates).

**Figure S15:**
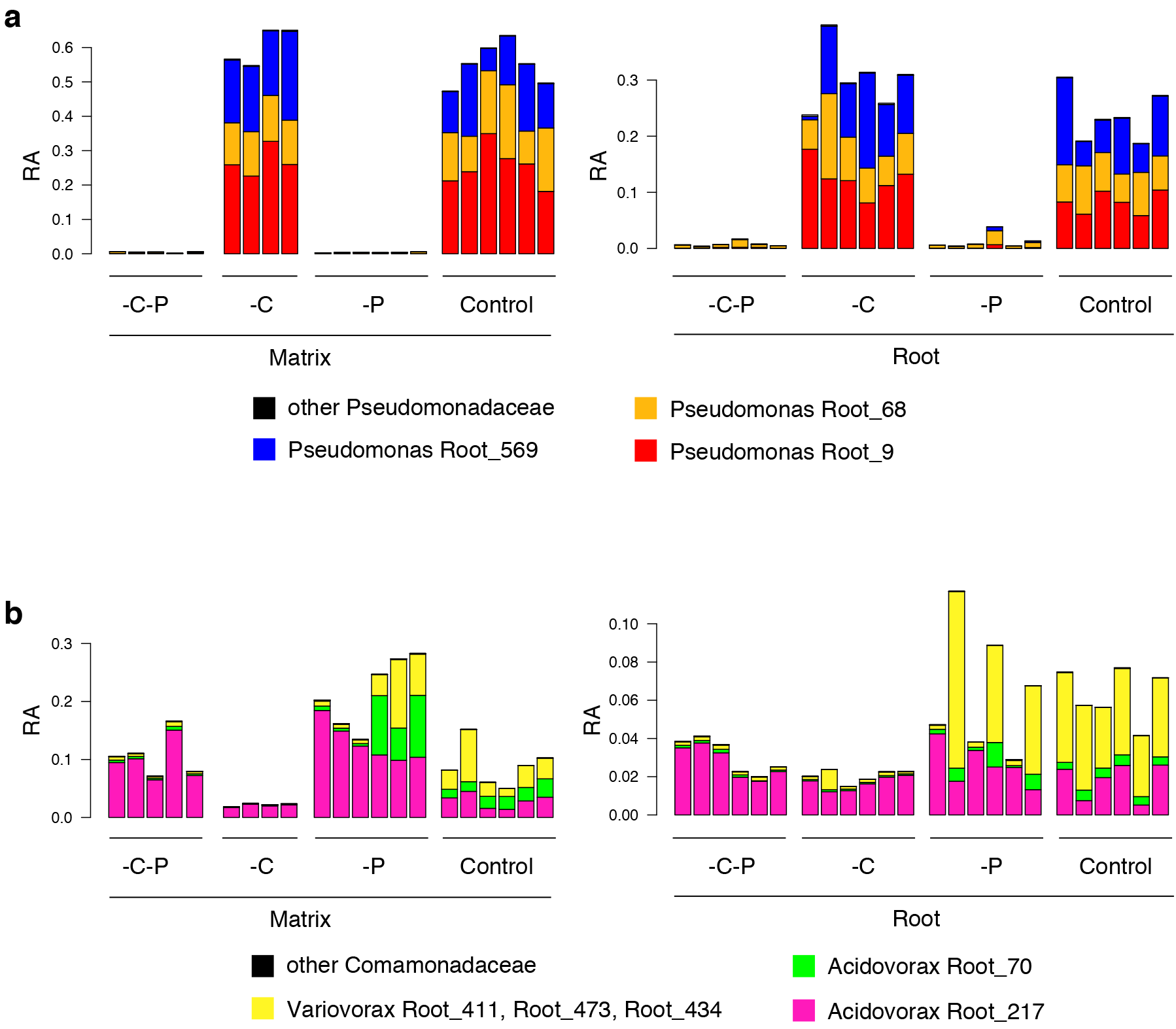
Validation of bacterial depletion in the FlowPot system. **a, b,** Relative abundances of isolates of the Pseudomonadaceae (**a**) and the Comamonadaceae (**b**) families in output matrix (left) and root (right) samples four weeks after inoculation in the FlowPot system. RA: relative abundance. Relative abundance of isolates belonging to Comamonadaceae (-C), Pseudomonadaceae (-P), Comamonadaceae and Pseudomonadaceae (-C-P) families is presented, together with the corresponding abundance in control samples inoculated with the full 148-member bacterial community.

**Figure S16:**
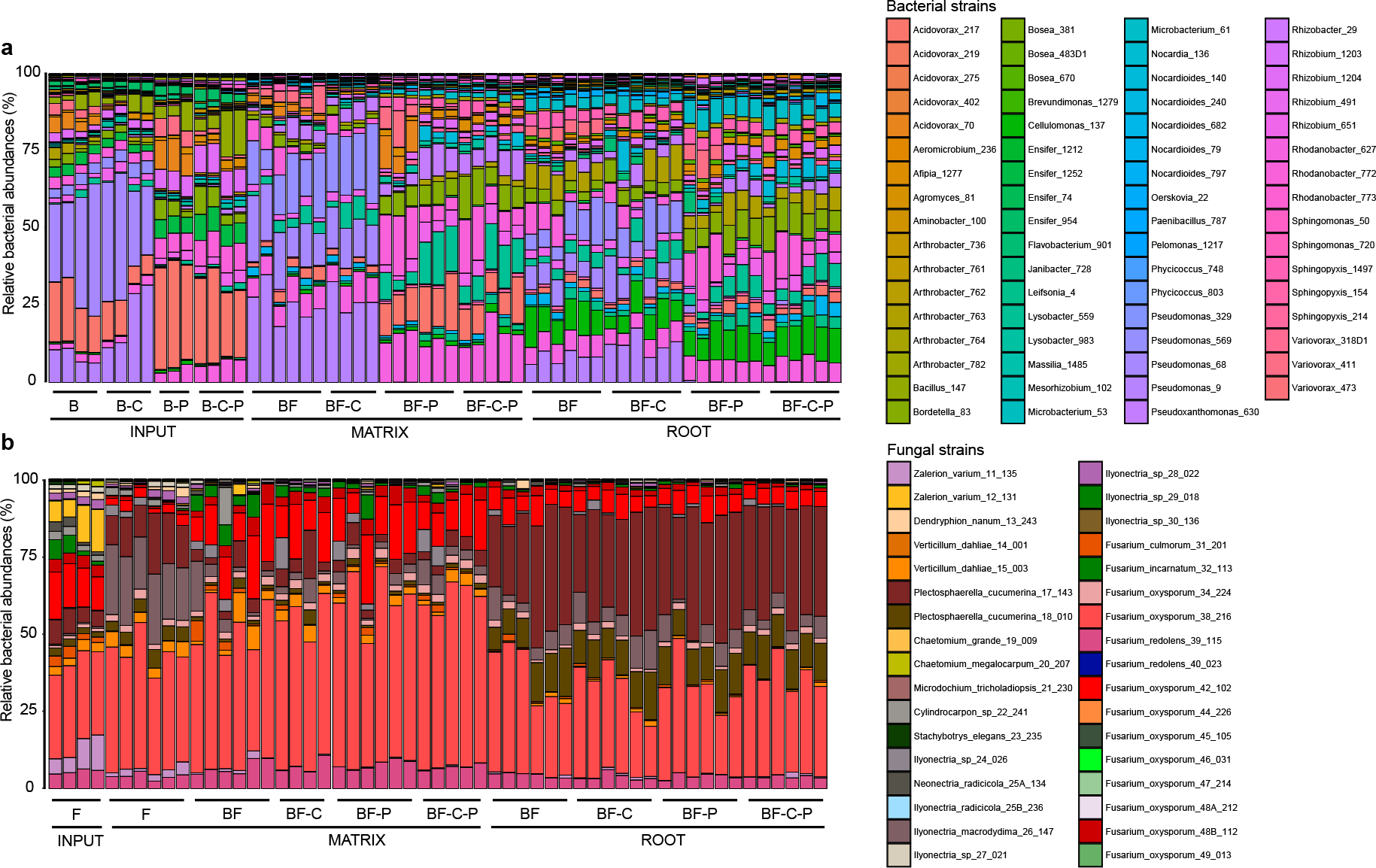
Abundance profiles of bacterial and fungal isolates in microbiota perturbation experiments. **a, b,** Relative abundances of bacterial (**a**) and fungal (**b**) strains in each microbial combination of the depletion experiment in input and output matrix and root samples four weeks after inoculation in the Flowpot system. Relative abundance of bacterial (**a**) and fungal (**b**) isolates (direct mapping at 100% sequence similarity) is presented for all conditions. B: full 148-member bacterial community. F: 34-member fungal community. -C: depletion of ten Comamonadaceae isolates. -P: depletion of eight Pseudomonadaceae isolates. -C-P: depletion of ten Comamonadaceae isolates and eight Pseudomonadaceae isolates.

**Figure S17:**
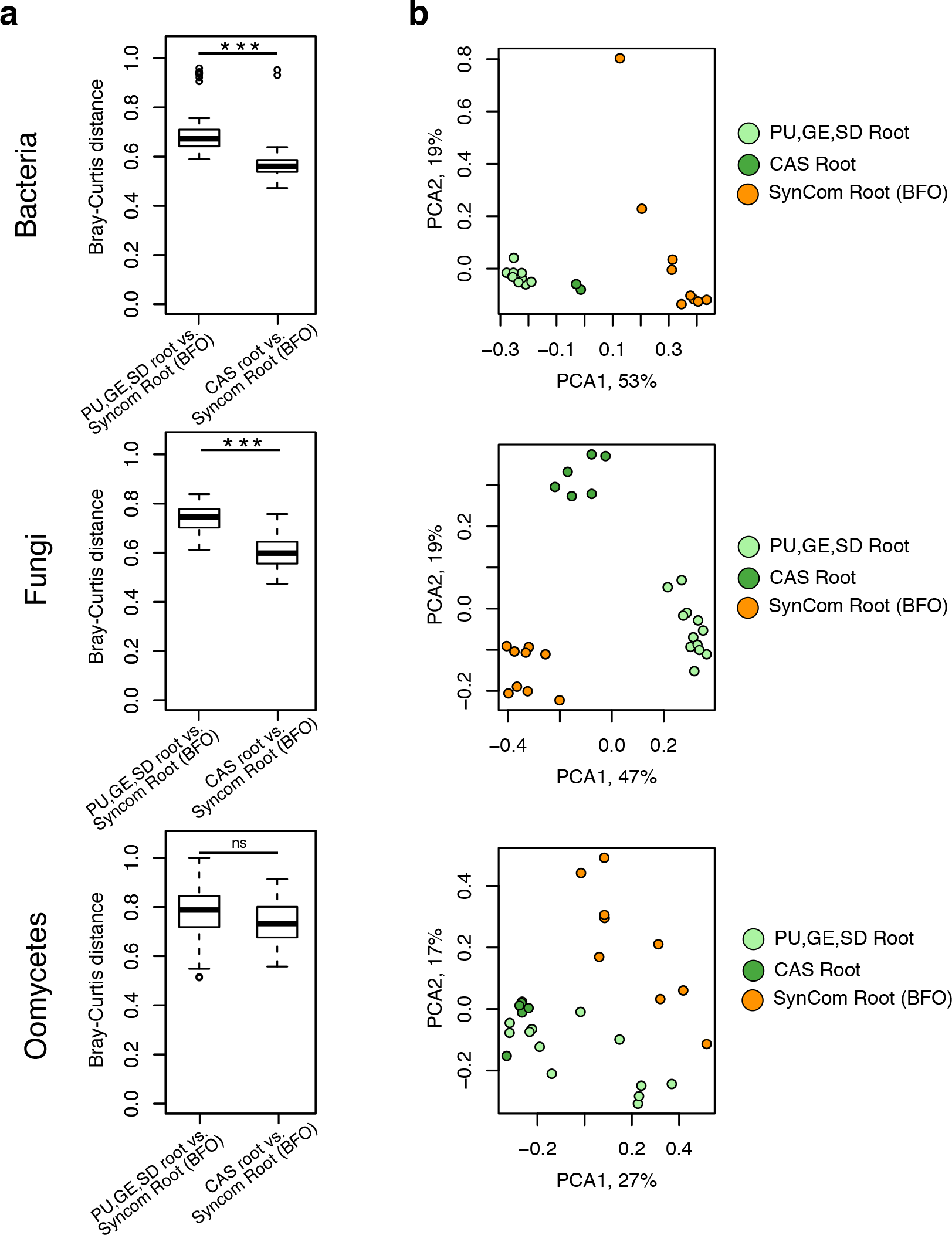
Comparison of the abundance of root-associated microbiome derived from natural sites and synthetic communities. **a,** Bray-Curtis distances between root samples from synthetic communities (SynComs) of the reconstitution experiment (see **Figure 4**) and root samples from one of the natural sites (PU, GE, SD) or from plants grown in Cologne agricultural soil (CAS). Kruskal-Wallis test, ns=not significant, *p<0.01,**p<0,001,***p<0.0001. **b,** Beta diversity determined using principal component analysis of the aforementioned root samples. For this analysis, only the 100 most abundant root-associated OTUs found in the three natural sites and in CAS samples were considered for calculation of distances between samples.

## Supplementary Tables

**Table S1: Location of the *A. thaliana* populations and corresponding soil characteristics**

**Table S2: Primers used in this study**

**Table S3: Root-derived fungal and oomycetal culture collections**

**Table S4: Microbial strains used for microbiota reconstitution experiments**

**Table S5: Contribution of different factors to microbial profile variance (PERMANOVA test, <0.05) and the incidence of specific microbial members**

**Table S6: Raw fluorescence data measured in the high-throughput fungal-bacterial interaction screen**

## References

Agler, M. T. et al. Microbial Hub Taxa Link Host and Abiotic Factors to Plant Microbiome Variation. PLoS Biol. 14, e1002352 (2016)

Almario, J. et al. Root-associated fungal microbiota of nonmycorrhizal Arabis alpina and its contribution to plant phosphorus nutrition. Proc. Natl Acad. Sci. USA 114, E9403–E9412 (2017)

Bai, Y. et al. Functional overlap of the Arabidopsis leaf and root microbiota. Nature 528, 364–369 (2015)

Bengtsson-Palme, J. et al. ITSx: Improved software detection and extraction of ITS1 and ITS2 from ribosomal ITS sequences of fungi and other eukaryotes for use in environmental sequencing. Methods Ecol Evol. 4, 914–919 (2013)

Boc, A., Boubacar Diallo, A. & Makarenkov, V. T-REX: a web server for inferring, validating and visualizing trees and networks. Nucleic Acids Res. 40, W573–579 (2012)

Bulgarelli, D. et al. Revealing structure and assembly cues for Arabidopsis root-inhabiting bacterial microbiota. Nature 488, 91–95 (2012)

Caporaso, J. G. et al. QIIME allows analysis of high-throughput community sequencing data. Nature Methods 7, 335–336 (2010)

Caporaso, J. G. et al. PyNAST: a flexible tool for aligning sequences to a template alignment. Bioinformatics 26, 266–267 (2009)

Castrillo, G. et al. Root microbiota drive direct integration of phosphate stress and immunity. Nature 543, 513–518 (2017)

Coleman-Derr, D., et al. Plant compartment and biogeography affect microbiome composition in cultivated and native Agave species. New Phytol. 209, 798–811 (2016)

Deshpande, V. et al. Fungal identification using a baysian classifier and the warcup training set of internal transcribed spacer sequences. Mycologia 108, 1–5 (2016)

DeSantis, T. Z. et al. Greengenes, a Chimera-Checked 16S rRNA gene database and workbench compatible with ARB. Appl. Environ. Microbiol. 72, 5069–5072 (2006)

Edgar RC. UPARSE: high accurate OTU sequences from microbial amplicon reads. Nature methods 10, 996–998 (2013)

Edwards, J, et al. Structure, variation, and assembly of the root-associated microbiomes of rice. Proc. Natl Acad. Sci. USA 112, E911–E920 (2015)

Faust, K. & Raes, J. CoNet app: inference of biological associations networks using Cytoscape. F1000Res. 5, 1519 (2016)

Figueroa-López, A. M., Cordero-Ramírez, J. D., Quiroz-Figueroa, F. R. & Maldonado-Mendoza, I. E. A high-throughput screening assay to identify bacterial antagonists against Fusarium verticillioides. J. Basic Microbiol. 54 Suppl 1, S125–S133 (2014)

Friedman, J. & Alm, E. J. Inferring correlation networks from genomic survey data. PLOS Comput Biol 8, e1002687012 (2012)

Garbaye, J. Helper Bacteria - a New Dimension to the Mycorrhizal Symbiosis. New Phytol. 128, 197–210 (1994)

Hacquard, S. et al. Microbiota and Host Nutrition across Plant and Animal Kingdoms. Cell Host Microbe 17, 603–616 (2015)

Hacquard, S. et al. Survival trade-offs in plant roots during colonization by closely related beneficial and pathogenic fungi. Nat. Commun. 7, 11362 (2016)

Hassani, M. A., Duran, P. B. & Hacquard, S. Microbial interactions within the plant holobiont. Microbiome 6, 58 (2018)

van der Heijden, M. G., de Bruin, S., Luckerhoff, L., van Logtestijn, R. S. & Schlaeppi, K. A widespread plant-fungal-bacterial symbiosis promotes plant biodiversity, plant nutrition and seedling recruitment. ISME J. 10, 389–399 (2016)

Hiruma, K. et al. Root Endophyte Colletotrichum tofieldiae Confers Plant Fitness Benefits that Are Phosphate Status Dependent. Cell 165, 464–474 (2016)

Katoh, K., Rozewicki, J. & Yamada, K. D. MAFFT online service: multiple sequence alignment, interactive sequence choice and visualization. Brief. Bioinform bbx108, (2017)

Keim, J., Mishra, B., Sharma, R., Ploch, S. & Thines, M. Root-associated fungi of Arabidopsis thaliana and Microthlaspi perfoliatum. Fungal Diversity 66, 99–111 (2014)

Kia, S. H. et al. Influence of phylogenetic conservatism and trait convergence on the interactions between fungal root endophytes and plants. ISME J. 11, 777–790 (2017)

Kremer, J.M., et al. FlowPot axenic plant growth system for microbiota research. BioRxiv https://doi.org/10.1101/254953 (2018)

Lancichinetti, A. & Fortunato, S. Community detection algorithms: A comparative analysis, Physcial review E 80, 056117 (2009)

Lebeis, S. L. et al. Salicylic acid modulates colonization of the root microbiome by specific bacterial taxa. Science 349, 860–864 (2015)

Lundberg, D. S. et al. Defining the core Arabidopsis thaliana root microbiome. Nature 488, 86–90 (2012)

Nilson, R. H. et al. A comprehensive, automatically updated fungal its sequence dataset for reference-based chimera control in environmental sequencing efforts. Microbes Environ. 30, 145–150 (2015)

Paulson, J. N., Stine, O. C., Corrado Bravo, H. & Pop, M. Differential abundance analysis in marker-gene surveys. Nature Methods 10, 1200–1202 (2013).

Ruggiero, M. A. et al. A Higher Level Classification of All Living Organisms. PLoS One 10:e0119248 (2015)

Santhanam, R., Weinhold, A., Goldberg, J., Oh, Y. & Baldwin, I. T. Native root-associated bacteria rescue a plant from a sudden-wilt disease that emerged during continuous cropping. Proc. Natl Acad. Sci. USA 112, E5013–E5020 (2015)

Shannon, P. et al. Cytoscape: a software environment for integrated models of biomolecular interaction networks. Genome Res. 13, 2498–2504 (2003)

Talbot, J. M. et al. Endemism and functional convergence across the North American soil mycobiome. Proc. Natl Acad. Sci. USA 111, 6341–6346 (2014)

Wang, Q., Garrity, G. M., Tiedje, J. M., & Cole, J. R. Naïve Baysian classifier for rapid assignment of rRNA sequences into the new bacterial taxonomy. Appl. Environm. Microbiol. 73, 5261–5267 (2007)

Zgadzaj, R. et al. Root nodule symbiosis in Lotus japonicas drives the establishment of distinctive rhizosphere, root and nodule bacterial communities. Proc. Natl Acad. Sci. USA 113, E7996–E8005 (2016)

Zhalnina, K. et al. Dynamic root exudate chemistry and microbial substrate preferences drive patterns in rhizosphere microbial community assembly. Nat. Microbiol. 3, 470–480 (2018)

